# Proteomic identification of *Coxiella burnetii* effector proteins targeted to the host cell mitochondria during infection

**DOI:** 10.1101/2020.05.28.121236

**Authors:** Laura F. Fielden, Nichollas E. Scott, Catherine S. Palmer, Chen Ai Khoo, Hayley J Newton, Diana Stojanovski

**Affiliations:** Department of Biochemistry and Molecular Biology and Bio21 Molecular Science and Biotechnology Institute, The University of Melbourne, Melbourne, VIC, Australia; Department of Microbiology and Immunology, University of Melbourne at the Peter Doherty Institute for Infection and Immunity, Melbourne, VIC, Australia

## Abstract

Modulation of the host cell is integral to the survival and replication of microbial pathogens. Several intracellular bacterial pathogens deliver a cohort of bacterial proteins, termed ‘effector proteins’ into the host cell during infection by sophisticated protein translocation systems which manipulate cellular processes and functions. Despite the importance of these proteins during infection the functional contribution of individual effectors is poorly characterised, particularly in intracellular bacterial pathogens with large effector protein repertoires. Technical caveats have limited the capacity to study these proteins during a native infection, with many effector proteins having only been demonstrated to be translocated during over-expression of tagged versions. Here we present development of a novel strategy to examine effector proteins in the context of infection. We coupled a broad, unbiased proteomics-based screen with organelle purification to study the host-pathogen interactions occurring between the host cell mitochondrion and the Gram-negative, Q fever pathogen *Coxiella burnetii.* We identify 4 novel mitochondrially-targeted *C. burnetii* effector proteins, renamed Mitochondrial *Coxiella* effector protein (Mce) B to E. Examination of the subcellular localisation of ectopically expressed proteins in epithelial cells confirmed the mitochondrial localisation, demonstrating the robustness of our approach. Subsequent biochemical analysis and affinity enrichment proteomics of one of these effector proteins, MceC, revealed the protein is imported into mitochondria and can interact with components of the mitochondrial quality control machinery. Our study adapts high-sensitivity proteomics to the study of intracellular host-pathogen interactions occurring during infection, providing a robust strategy to examine the sub-cellular localisation of effector proteins during native infection. This approach could be applied to a range of pathogens and host cell compartments to provide a rich map of effector dynamics throughout infection.

## Introduction

Numerous microbial pathogens have evolved strategies to survive within the host cell. During infection, a subset of bacterial virulence factors, termed effector proteins, can be delivered into the host cell where they function to modulate host cell processes and ultimately contribute to bacterial pathogenesis. Bacterial effector proteins are directly translocated into the host cell by specialised secretion systems (*1, 2*). The repertoire of effector proteins encoded by a bacterial pathogen is unique, reflective of the intracellular niche it occupies and varies greatly across different bacterial species (*3, 4*). Bacteria possessing a Dot/Icm type 4 secretion system (hereafter referred to as T4SS) typically encode a large effector cohort, for instance the respiratory pathogen *Legionella pneumophila* encodes for approximately 330 effector proteins and the evolutionarily related, *Coxiella burnetii* for approximately 150 (*5, 6*). The substantial number of proteins potentially delivered by these bacterial pathogens into the host cell creates a problem in identifying *bona fide* translocated effectors and delineating the function of these proteins during infection.

Approaches to biochemically characterising effector proteins typically concentrate on one effector in isolation, often removed from the context of infection. The low abundance of effector proteins within the host cell creates a technical challenge that has limited study during native infection. Often, over-expression or ‘tagging’ of effectors with a protein or peptide label is employed to enable localisation and interaction studies. However, our increasing understanding of the manner in which bacterial secretion systems deliver protein substrates into the host cell and the important role of meta-effector interactions during infection highlights the risks associated with an individualistic approach (*7-11*). The rapidly expanding field of high-sensitivity mass-spectrometry coupled to sub-cellular organelle isolation presents a solution to the study of endogenous host-pathogen interactions occurring during infection, particularly for bacterial pathogens harbouring large effector repertoires where functional redundancy exists within the effector cohort.

*C. burnetii* is a Gram-negative, obligate intracellular bacterial pathogen and the causative agent of the zoonotic disease Query (Q) fever in humans (*12*). Q fever is a complex disease with a range of clinical presentations from asymptomatic seroconversion to acute illness and debilitating chronic infection (*13-16*). Human infection occurs by inhalation of contaminated aerosols and the bacterium preferentially infects alveolar macrophages (*17*). *C. burnetii* encodes a T4SS which is essential for intracellular replication of the bacterium (*18, 19*). This translocation system mediates the delivery of bacterial effector proteins into the host cell over the course of infection (*20, 21*). Thus far, 150 proteins have been identified with the capacity for T4SS translocation, however, whether these are all delivered into the host cell during native infection is unknown. The identification and characterisation of substrates of the *C. burnetii* T4SS has been a rapidly expanding area of research, greatly assisted by the development of axenic culture and genetic manipulation techniques (*18, 22-27*). Despite significant advancement in our identification of these bacterial virulence factors, the majority of *C. burnetii* effector proteins remain functionally uncharacterised.

We previously demonstrated that the *C. burnetii* effector protein MceA specifically targets the host cell mitochondria during infection (*28*). This research highlighted the organelle as a *bona fide* target of *C. burnetii* during infection. Mitochondria are essential eukaryotic organelles, fundamental to cell function and survival. Mitochondria contribute to metabolism and bioenergetics, iron-sulfur cluster biogenesis, lipid synthesis, calcium homeostasis, immune signalling and apoptosis (*29-31*). Given this crucial role in cell biology and the important involvement of mitochondria in the host response to infection, mitochondrial functions are frequently targeted by microbial pathogens during infection of the host cell (*32-34*). For instance, *L. pneumophila* targets mitochondrial dynamics and metabolism via the T4SS effector protein MitF (*35*). MitF targets host cell Ran GTPase promoting DRP1- dependent fragmentation of the host mitochondria and assisting in biphasic regulation of mitochondrial respiration in a macrophage host (*35*).

Currently, little is known about the interaction between *C. burnetii* and the host cell mitochondrion during infection. Earlier studies demonstrated that the bacterium has the ability to block apoptosis by preventing the release of cytochrome *c* from the mitochondria (*36*). The ability to inhibit the mitochondrial pathway of apoptosis has highlighted two additional C. *burnetii* effector proteins, AnkG and CaeB, that appear to target the organelle however, characterisation of these effectors was performed in the absence of infection (*37, 38*). In consideration of the large cohort of bacterial proteins delivered to the host cell by *C. burnetii* and the essentiality of multiple mitochondrial functions to cellular homeostasis, we hypothesised that additional effector proteins would localise to the organelle during infection.

Here, an unbiased, unlabelled proteomics approach was utilised to identify *C. burnetii* effector proteins associated with the host cell mitochondria during infection. The application of this approach has established a list of 7 candidate *C. burnetii* proteins targeted to the host cell mitochondria and validated the association of 4 proteins with the organelle which we renamed MceB - E. We further characterise one of these candidates, CBU1425 (later renamed MceC), for confirmation of submitochondrial localisation and protein interactions. MceC was found to be imported into the organelle and likely integrated into the inner membrane. Furthermore, we demonstrate that MceC forms associations with members of a mitochondrial inner membrane quality control network. Development and use of this unbiased proteomics-based technique has allowed us to uncover host-pathogen interactions between *C. burnetii* and the mitochondria in the context of native infection and provides an exciting platform for future exploration of effector interactions with host cell organelles.

## Materials and methods

### Bacterial strains and culturing conditions

Plaque purified *Coxiella burnetii* Nine Mile Phase II (NMII), strain RSA439 clone 4 wildtype and wildtype expressing mCherry were used in this study. *C. burnetii* strains were cultured axenically in liquid Acidified Citrate Cysteine Medium 2 (ACCM-2) with chloramphenicol (3 μg/ml, Boehringer Mannheim) when required for 7 days at 37 °C, 5% CO_2_ and 2.5% O_2_, as previously described (*22, 39*). *Escherichia coli* XL1-Blue, used for plasmid construction and propagation were cultured in Luria-Bertani (LB) broth or agar plates containing ampicillin (50 μg/ml, Sigma) as appropriate.

### Mammalian cell lines and culturing, transient transfection and stable cell line generation

Cell lines used in this study were THP-1 (human monocytic leukemia) cells, HeLa (Henrietta Lacks, human cervical carcinoma cells, CCL2) and HEK293T Flp-In T-REx 293 (Thermo Fisher Scientific). THP-1 cells were cultured in Roswell Park Memorial Institute (RPMI) 1640 medium containing 10% heat-inactivated Fetal Bovine Serum FBS (FBS; Thermo Fisher Scientific). HeLa and HEK293 cells were cultured in Dulbecco’s Modified Eagle Medium (DMEM, Gibco) containing 5 - 10% FBS (Thermo Fisher Scientific). All cells were maintained at 37 °C, 5% CO_2_.

Transient transfections were performed using Lipofectamine® 3000 (Thermo Fisher Scientific) transfection reagent as per the manufacturer’s instructions. Cells were incubated with transfection media for 6 - 8 hours prior to replacement with fresh DMEM containing 5% FBS and incubated at 37 °C for appropriate time (typically a total of 18 hours unless otherwise stated).

Stable tetracycline-inducible cell lines were generated in accordance with the manufacturers protocol and as previously described (*40*). Briefly, cells were seeded into a 6-well plate to reach a confluency of ˜60% on the day of transfection. Cells were co-transfected with pcDNA5-FRT/TO-CBU1425- 3XFLAG and pOG44 (encoding the Flp-recombinase) at a 1:9 ratio (per ng of DNA). Three days post-transfection, positive clones were selected for using Hygromycin B (200 μg/ml; Thermo Fisher Scientific). Selection was applied until single colonies could be detected and foci resuspended and transferred for recovery and expansion. Protein expression was induced by the addition of tetracycline (1 μg/ml; Sigma) for desired time (determined during time course).

#### C. burnetii *quantitation and infections*

THP-1 cells were seeded at a density of 1.5 × 10^7^ cells per well into 15 cm tissue culture plates and treated with 10 nM phorbol 12-myristate 13-acetate for 3 days to induce differentiation into a macrophage-like cell. *C. burnetii* quantitation was performed using quantitative PCR (qPCR) with gene specific primers for *ompA* (Forward: 5’-CAGAGCCGGGAGTCAAGCT-3’, Reverse: 5’- CTGAGTAGGAGATTTGAATCGC-3’) to provide a multiplicity of infection (MOI) of 100 (*41*). Cells were incubated with bacteria at 37 °C, 5% CO_2_ for 24 hours, washed once in warm 1X PBS, media replaced with fresh RPMI 1640 with 10 *%* FCS and incubated for a further 48 hours prior to collection for mitochondrial isolation and proteomics analysis.

### Immunofluorescence microscopy

Cells were fixed in paraformaldehyde (PFA: 4% (w/v) in PBS containing 5% (w/v) sucrose) for 10 - 15 min and then permeabilised in 0.1% (v/v) TX-100 in PBS at RT. Coverslips were blocked in 3% (w/v) BSA in PBS for 30 min at RT, followed by incubation with primary and secondary antibodies diluted in 3% (w/v) BSA in PBS. Coverslips were incubated with primary antibodies for 1 - 2 hours at RT, washed with PBS (3 changes over 10 min) followed by incubation with secondary antibodies for 30 - 60 min (1:500 dilution; goat anti-mouse AlexaFluor 488 conjugate (Invitrogen) or goat anti-rabbit AlexaFluor 568 conjugate (Invitrogen)). Coverslips were washed with PBS containing Hoechst stain (10 μg/mL; Invitrogen), affixed onto glass slides using mounting media (0.2 M DABCO (Sigma), 0.1 M Tris-HCl (pH 8.0), 90% glycerol) and sealed using nail polish. Slides were imaged on a Leica SP8 confocal microscope and image analysis performed using ImageJ/FIJI software (https://imagej.nih.gov/ij/).

### Mitochondrial isolation and purification

Mitochondria were isolated from tissue culture cells by differential centrifugation as previously described (*42*). Isolated cells were resuspended in solution A (70 mM sucrose, 220 mM mannitol, 20 mM HEPES-KOH (pH 7.6), 1 mM EDTA, 0.1mM PMSF and 2 mg/mL BSA) and homogenised in a handheld glass Dounce homogeniser (typically 15 - 20 strokes). Cellular homogenate was centrifuged at 600 *g* for 5 min at 4 °C to remove nuclear and cellular debris. The supernatant was centrifuged at 12 000 *g* for 10 min at 4 °C following which the pellet (crude mitochondrial fraction) was resuspended in solution B (70 mM sucrose, 220 mM mannitol, 20 mM HEPES-KOH (pH 7.6) and 1 mM EDTA). Mitochondria isolated from *C. burnetii-infected* cells were further purified using sucrose density gradient centrifugation and affinity purification. Sucrose density gradient centrifugation was performed as previously described with modifications (*43*). Following crude isolation, mitochondria were resuspended in continuous gradient buffer (0.25 M sucrose, 10 mM Tris-Cl (pH 7.4) and 1 mM EDTA) and overlaid on a 34 - 64% continuous sucrose gradient with a 67% sucrose cushion at the base of the tube (34/64/67% (w/v) sucrose, 10 mM Tris-Cl (pH 7.4) and 1 mM EDTA). Gradients were centrifuged at 170 000 *g* for 1 hour in a SW41Ti swinging bucket rotor (Beckman Coulter) at 4 °C. Following ultracentrifugation, 12 × 1 ml fractions were removed and fractions 2 - 5 were pooled, diluted 1:2 with continuous gradient buffer and centrifuged at 16 000 *g* for 10 mins at 4 °C to isolate mitochondria. Mitochondria were further purified by affinity purification using anti-TOM22 microbeads (Miltenyi Biotec) as per the manufacturers protocol with several modifications. Briefly, mitochondria were resuspended in 1 ml solution B containing protease inhibitor (PI; Roche) and incubated with anti-TOM22 microbeads on a rotary wheel for 30 mins at 4 °C. The mitochondrial resuspension was transferred to a MACS LS column (Miltenyi Biotech) and bound mitochondria were washed 3X with solution B containing PI and eluted in solution B containing PI. Purified mitochondria were re-isolated and prepared for analysis by mass spectrometry. Mitochondrial protein concentration was estimated using UV spectrophotometry or BCA protein assay (Pierce).

### Gel electrophoresis and immunoblot analysis

Tris-tricine SDS-PAGE was performed as previously described (*44-47*). Samples for electrophoresis were combined with SDS-PAGE loading dye (50 mM Tris-Cl (pH 8.45), 100 mM dithiothreitol, 2% [w/v] SDS, 10% [v/v] glycerol, 0.1% [w/v] bromophenol blue). Electrophoresis was performed in the presence of Tris-tricine SDS -PAGE cathode buffer (0.1 M Tris, 0.1 M Tricine (pH 8.45), 0.1% [w/v] SDS) and anode buffer (0.2 M Tris-Cl (pH 8.9).

Western transfers were performed using the OWI^™^ HEP-1 Semidry Electroblotting System (ThermoFisher Scientific) semi-dry transfer method. Following electrophoresis SDS-PAGE gels were transferred onto a polyvinylidene fluoride (PVDF) membrane (Millipore). PVDF membranes (whole or in strips) were subjected to immunoblot analysis with specific primary antibodies (*Table 1*) and secondary antibodies (anti-mouse/rabbit IgG coupled to horseradish peroxidase; Sigma Aldrich; 1:5000 dilution in blocking buffer). Detection of chemi-luminescent signal was performed with Clarity ECL Western Blotting substrate (BioRad) and imaged using a ChemiDoc^™^ MP Imaging system (BioRad).

### Mitochondrial treatments

Mitochondrial subfractionation were performed with mitochondria isolated by differential centrifugation. Samples (100 μg) were resuspended in: (i) solution B, (ii) swelling buffer (10 mM HEPES-KOH, (pH 7.4)) or (iii) solubilisation buffer (1% [v/v] TX-100) and incubated on ice for 5 min. Samples were incubated for a further 10 min in either the absence or the presence of Proteinase K (PK) (50 μg/ml) followed by addition of 1mM PMSF for 5 - 10 min. All samples were 12.5% (w/v) TCA precipitated at 4 °C overnight and analysed by SDS-PAGE and immunoblotting. For sodium carbonate extraction, isolated mitochondria (200 μg) were resuspended in 100 mM Na_2_CO_3_ (pH 11) and incubated on ice for 30 min with occasional agitation. Samples were centrifuged at 100 000 *g* for 30 min at 4 °C to separate integral membrane proteins (pellet fraction) and soluble proteins (supernatant fraction). Both fractions were TCA precipitated and analysed by SDS-PAGE and immunoblotting.

### Immunoprecipitation analysis

Isolated crude mitochondrial samples (3.5 - 5 mg) were resuspended in digitonin solubilisation buffer (1% [v/v] digitonin, 50 mM Tris-Cl (pH 7.4), 150 mM NaCl) at 2 mg/ml and incubated, end-over-end at 4 °C for 30 min. Solubilised mitochondria were clarified by centrifugation at 16 000 *g* for 10 min at 4 °C. The supernatant was diluted to a final detergent concentration of 0.1% digitonin and incubated with pre-equilibrated anti-FLAG resin (Sigma Aldrich- A2220; 5 - 10 μl of resin/mg mitochondrial protein), end-over-end for 30 min at 4 °C. Resin and bound proteins were transferred to a Pierce spin column (Thermo Fisher Scientific-69705) and washed 8X in solubilisation buffer containing 0.1% [w/v] digitonin. Bound proteins were eluted in 0.2 M glycine (pH 2.5) over 2 by 5 min incubations and elution fractions pooled. Total, supernatant, unbound, wash and elution fractions were TCA precipitated and pellet fraction directly resuspended into SDS-PAGE loading dye. All fractions were analysed by SDS-PAGE and immunoblotting, or elution fraction prepared for analysis by mass spectrometry.

### Mass spectrometry

Mitochondrial protein (50-100 μg) was acetone-precipitated (8 volumes acetone, 1 volume water, 1 volume sample) overnight at −20 °C. Precipitated material was then spun down at 16 000 *g* for 10 mins at 4 °C and dried to remove remaining acetone at 65 °C for 5 - 10 mins. Pellets were resuspended in Urea (6M, Sigma), Thiourea (2M, Sigma) and DTT (10 mM, Austral Scientific) and incubated at RT in the dark for 1 hour to reduce disulfide bonds. Samples were then alkylated by the addition of chloroacetamide (CAA; 40 mM) and incubated for a further hour. Alkylation was then halted by the addition of DTT (50 mM) and incubation at RT for 15 min. Samples were digested with Lys-C (1/200 (w/w); Wako Lab Chemicals) for 4 hours at RT before dilution with ammonium bicarbonate (20 mM; Sigma) and digestion with trypsin (1/50 (w/w); Sigma) overnight at 25 °C. Digested peptides were then acidified by the addition of formic acid (2% (v/v); Sigma) and desalted using homemade C18 stage tips (Empore C18, Sigma) before being dried down for analysis by liquid chromatography mass spectrometry (LC-MS) (*48, 49*). Prior to loading, samples were reconstituted in MS running buffer (2% acetonitrile (ACN, Sigma), 0.1% trifluoroacetic acid (TFA, Sigma)) to a concentration of 0.5 μg/μl of which 2 μg was loaded for mass spectrometry.

Label-free quantitative (LFQ) mass spectrometry was performed to identify *C. burnetii* proteins co-purifying with mitochondria. Mitochondria isolated from *C. burnetii*_mCherry_-infected THP-1 cells were analysed by LC-MS on an Orbitrap Lumos^™^ mass spectrometer (ThermoFisher Scientific) coupled to a Dionex Ultimate 3000 Ultra-Performance Liquid Chromatography (ThermoFisher Scientific) using a two-column chromatography set up composed of a PepMap100 C18 20 mm × 75 μm trap and a PepMap C18 500 mm × 75 μm analytical column (Thermo Fisher Scientific). Samples were concentrated at 5 μl/min onto the trap column for 5 minutes before the trap column was switched in-line with the analytical column using Buffer A (0.1% Formic acid, 5% DMSO). Analytical separation was performed at 300 nl/min using a non-linear ACN gradient by altering the composition of Buffer B (0.1% (v/v) formic acid, 94.9% (v/v) ACN, 5% (v/v) DMSO) from 3% to 22% over 180 min then from 30% to 40% over 7 min, 40% to 90% over 5 min, holding at 90% for 5 minutes then dropped to 3% Buffer B over 3 min with the column then equilibrated by holding at 3% Buffer B for 5 min. Data were collected in positive mode using Data Dependent Acquisition with a 120 000 orbitrap resolution MS1 scan of mass-to-charge (m/z) range of 375 - 1500 (automatic gain control (AGC) set to 4 × 10^5^ or a maximum injection time of 50 ms) acquired every 3 sec followed by MS2 scans. MS2 scans were acquired using high-energy collision dissociation fragmentation with a normalised collision energy of 35%, with an isolation window of 1.6 in the quadrupole, resolution of 7500 and an AGC of 5 × 10^4^ or a maximum injection time of 22 ms. A dynamic exclusion of duration of 30 sec was applied for repeated precursors. Raw files were analysed using MaxQuant platform (version 1.6.2.10) and searched against Uniprot human database (Accession: UP000005640; downloaded October 2018) and *C. burnetii* database (Accession: UP000002671; downloaded 2017), containing reviewed, canonical and isoform variants and a database containing common contaminants generated by the Andromeda search engine (*50*). LFQ search parameters were left as default with Trypsin/P specificity with a maximum of 2 missed cleavages. Searches were performed with cysteine carbamidomethylation as a fixed modification and methionine oxidation, N-terminal acetylation and methionine oxidation as variable modifications. False discovery rate (FDR) was determined using the target-decoy approach set to 2% for peptides and 1% for proteins. Unique and razor peptides were used for identification with a minimum ratio count of 2. A search tolerance of 4.5 ppm was used for MS1 and 20 ppm for MS2 matching. ‘Re-quantify’ and ‘match between runs’ functions were enabled with a match time window of 0.7 min (*51*).

Samples prepared following immunoprecipitation from HEK293 MceC^3XFLAG^ cell line were analysed by LC-MS on an Orbitrap Elite^™^ mass spectrometer (ThermoFisher Scientific) coupled to a Dionex Ultimate 3000 Ultra-Performance Liquid Chromatography (ThermoFisher Scientific) using a two-column chromatography set up composed of a PepMap100 C18 20 mm × 75 μm trap and a PepMap C18 500 mm × 75 μm analytical column (Thermo Fisher Scientific). Samples were concentrated at 5 μl/min onto the trap column for 5 minutes. Analytical separation was performed at 300 nl/min using a 65 min non-linear gradient by altering the concentration of Buffer B (5% (v/v) DMSO, 94.9% (v/v) ACN and 0.1% (v/v) formic acid) from 0 to 3% over 5 mins, 3% to 22% over 32 min, then from 22% to 40% over 10 min, 40 to 80% over 5 min and held at 80% for 5 min then dropped to 3% over 3 min with the column equilibrated by holding at 3% for 10 min. Data were collected in positive mode using Data Dependent Acquisition with a 100 000 orbitrap resolution MS1 scan of mass-to-charge (m/z) range of 300 - 1650. The top 20 most intense precursor ions were subjected to rapid collision induced dissociation with normalised collision energy of 30 and activation q of 0.25. A dynamic exclusion of duration of 30 sec was applied for repeated precursors. Raw files were analysed using MaxQuant platform (version 1.6.10.43) and searched against Uniprot human database (October 2018) and fasta sequence for the ORF of CBU1425 (downloaded from Uniprot October 2019). LFQ search parameters were left as default with Trypsin/P specificity with a maximum of 2 missed cleavages. Oxidation of methionine and N-terminal acetylation were specified as variable modifications. Carbamidomethylation of cysteine was set as a fixed modification. A search tolerance of 4.5 ppm was used for MS1 and 0.5 Da for MS2 matching. FDR were determined using the default target-decoy approach set to 1% for both peptides and proteins. Match between runs was enabled with a match time window of 0.7 min.

Mitochondria isolated from HEK293 MceC_3XFLAG_ or empty vector (EV) cell lines during expression timecourse were analysed on a Q Exactive PIUS^™^ orbitrap mass spectrometer (ThermoFisher Scientific) coupled to a Dionex Ultimate 3000 Ultra-Performance Liquid Chromatography (ThermoFisher Scientific). Peptides were injected into the trap column (PepMap100 C18 20 mm × 100 μm; ThermeFisher Scientific) at a flow rate of 5 μl/min before changing the trap in-line with the analytical column (PepMap C18 500 mm × 75 μm; ThermoFisher Scientific). Analytical separation was performed at 300 nl/min non-linear gradient by altering the composition of Buffer B from 3% to 22% over 105 min then from 30% to 40% over 7 min, 40% to 90% over 5 min, holding at 90% for 5 minutes then dropped to 3% Buffer B over 3 min with the column then equilibrated by holding at 3% Buffer B for 5 min) over 130 min.

Data were collected in positive mode using Data Dependent Acquisition with a 70 000 orbitrap resolution MS1 scan of mass-to-charge (m/z) range of 375 - 1400 (maximum injection time of 50 ms, AGC target of 3 × 10^6^). The top 15 most intense precursor ions were subjected to MS2 using high-energy collision dissociation fragmentation with a normalised collision energy of 28% (resolution of 17 500, AGC target of 5 × 10^4^ and maximum injection time of 50 ms). Raw files were analysed using MaxQuant platform (version 1.6.5.0) and searched against Uniprot human database (October 2018) and fasta sequence for the ORF of CBU1425 (downloaded from Uniprot October 2019). LFQ search parameters were left as default with Trypsin/P specificity with a maximum of 2 missed cleavages. Oxidation of methionine and N-terminal acetylation were specified as variable modifications. Carbamidomethylation of cysteine was set as a fixed modification. A search tolerance of 4.5 ppm was used for MS1 and 20 ppm for MS2 matching. FDR were determined using the default target-decoy approach set to 1% for both peptides and proteins. Match between runs was enabled with a match time window of 0.7 min.

### Data analysis in Perseus

The ‘protein groups’ output file was imported into the Perseus platform for further processing (*52*). Identifications labelled by MaxQuant as ‘only identified by site’, ‘potential contaminant’ and ‘reverse hit’ were removed and LFQ values normalized by log_2_-transformation. Identifications were matched to human and *C. burnetii* annotation files (downloaded from Uniprot at the same time as databases) using gene name identifiers. Mitochondrial proteins were annotated using the human MitoCarta database (*53, 54*). Data was exported into Excel and then into the R framework (https://www.r-project.org) integrated with R studio (https://rstudio.com/products/rstudio/) for generation for generation of graphics.

### Data availability

The mass spectrometry proteomics data have been deposited to the ProteomeXchange Consortium via the PRIDE (*55, 56*) partner repository with the dataset identifier PXD019322 (mitochondrial purification from *C. burnetii*-infected cells), PXD019417 (MceC affinity purification and interactome analysis) and PXD019337 (Mitochondrial proteome analysis during MceC expression time-course).

### Bioinformatic analysis of *C. burnetii* proteins using S4TE

*C. burnetii* proteins were analysed and scored based on probability of translocation via the T4SS using the online bioinformatics program Searching Algorithm for Type IV Effector proteins (S4TE) 2.0, (https://sate.cirad.fr) (*57*). Pre-loaded *C. burnetii* Nine Mile RSA493 chromosome and plasmid (pQpH1) genbank sequences were analysed using default settings (https://sate.cirad.fr/S4TE.php). Data were exported to Microsoft Excel and unannotated effector proteins manually annotated. Data were imported into the R framework (https://www.r-project.org) integrated with R studio (https://rstudio.com/products/rstudio/) for generation for comparison with *C. burnetii* protein enrichment alongside purified mitochondria and generation of graphics.

### Experimental design and statistical rationale

For label free quantitative LC-MS/MS experiments of mitochondria, statistical analysis of data was consistent with published analyses from our and other groups employing similar instrumentation and methods (*46, 62, 63*). All experiments were performed in at least 3 independent biological replicates. Data was imported into Perseus analysis platform and LFQ values log_2_ transformed. Enrichment ratio was calculated on normalised data, requiring a minimum of 2 valid values in either ‘crude’ or ‘pure’ mitochondria group. Imputation (downshifted by 1.8σ with a distribution of 0.3σ) was applied to missing values in both groups population missing datapoints with values equivalent to the limit of detection in each experiment. Individual replicate enrichment ratios were averaged and compared to S4TE T4SS translocation probability scores. Proteins with an enrichment ratio greater than 0.85 and translocation score greater than 72 were considered highly enriched at the mitochondria. MceC affinity enrichment analysis were performed on mitochondria isolated from 3 biological replicates for each control and MceC-expressing cells requiring a minimum of 3 valid values in replicates from MceC-expressed condition. The fold change value used for significance in MceC affinity enrichment analysis was determined through a two-sided t-test based with multiple hypothesis undertaken using a permutation-based FDR with an FDR of 5% and s0 value of 1. Imputation (downshifted by 1.8σ with a distribution of 0.3σ) was applied to missing values. Label-free MceC time course experiments were performed on 3 biological replicates per time point. Imputation (downshifted by 1.8σ with a distribution of 0.3σ) was applied to missing values in both groups population missing datapoints with values equivalent to the limit of detection in each experiment. An analysis of variance (ANOVA) test with multiple hypothesis using a permutation-based FDR with an of 5% and an s0 value of 0 was used to identify proteins significantly altered across all time-points.

### Antibodies

**Table 1:** Antibodies used in this study.

| Antibody | Source | Identifier | Dilution |
| --- | --- | --- | --- |
| Mouse monoclonal FLAG | Sigma-Aldrich | Catalogue #: F1804 | WB: 1:2000<br>IFA: 1:200 |
| Rabbit polyclonal mCherry | Novus Biologicals | Catalogue #: NBP2-25157 | WB: 1:1000 |
| Mouse monoclonal β-actin | Sigma-Aldrich | Catalogue #: A5316 | WB: 1:5000 |
| Mouse monoclonal Cytochrome c | BD Biosciences | Catalogue #:556433 | WB: 1:300 |
| Rabbit monoclonal Histone H3 | Cell Signalling | Catalogue #: 4499 | WB: 1:2000 |
| Rabbit polyclonal Tim44 | Proteintech | Catalogue #: 138-59-1-AP | WB: 1:500 |
| Rabbit polyclonal Tim29 | Sigma-Aldrich | Catalogue #: T8954 | WB: 1:1000 |
| Mouse monoclonal SDHA | Abcam | Catalogue #: ab14715 | WB: 1:1000 |
| Mouse monoclonal LAMP1 | DSHB | Catalogue #: H4A3-C | WB: 1:1000<br>IFA: 1:250 |
| Mouse monoclonal PDI | Sapphire Bioscience | Catalogue #: ADI-SPA-891-D | WB: 1:500 |
| Mouse monoclonal Pex14 | Proteintech | Catalogue #: 10594-1-AP | WB:1:1000 |
| Mouse monoclonal OPA1 | BD Biosciences | Catalogue #: 612606 | WB: 1:1000 |
| Rabbit polyclonal Bak | The Ryan Laboratory | (58) | WB: 1:500 |
| Rabbit polyclonal NDUFAF2 | The Ryan Laboratory | (59) | WB: 1:500<br>IFA: 1:350 |
| Rabbit polyclonal NDUFA9 | The Ryan Laboratory | (60) | WB: 1:1000 |
| Rabbit polyclonal Mfn2 | The Ryan Laboratory | (61) | WB: 1:500 |

## Results

### Purification of mitochondria from *C. burnetii*-infected cells

In order to identify effector proteins targeted to the mitochondria during *C. burnetii* infection an unbiased, unlabelled, proteomic approach was employed. An experimental pipeline to obtain pure mitochondria from *C. burnetii* infected cells for downstream proteomic analysis was required. The purity of this sample was vital to aid in the identification of high-confidence mitochondrial associated effector proteins, as contamination with *C. burnetii* would result in the misidentification of translocated proteins. A further aspect to consider was contamination of mitochondria with other host cell organelles, which could decrease the mitochondrial specificity of the identified effector proteins. A standard mitochondrial isolation, using differential centrifugation, yields ‘crude’ mitochondrial samples with a low degree of purity hence we opted for a more stringent approach based on published methods to purify mitochondria from complex mammalian cell lysates (*43, 64-67*). As these published methods had not been performed in the context of bacterial infection we therefore asked if one, or a combination, of these approaches could be applied to purify mitochondria from *C. burnetii*-infected cells.

Mitochondria were isolated by differential centrifugation from HeLa cells persistently infected with a strain of *C. burnetii* constitutively expressing mCherry (*C. burnetii*_mCherry_). Expression of mCherry by *C. burnetii* functioned as a downstream marker for bacterial detection. This crude mitochondrial lysate was overlaid on a 34 - 64% (w/v) continuous sucrose gradient and separated by ultracentrifugation *(Figure 1A*). Mitochondria previously isolated by this method were recovered at a sucrose density of ˜40% (w/v) (*43*). Fractions were collected and proteins separated by SDS-PAGE to analyse for organelle and bacterial content (*Figure 1B*). Interestingly, mitochondria and *C. burnetii* appeared to separate into distinct fractions of the continuous gradient *(Figure 1B, ‘mitochondria’ lanes 3 - 6 and* ‘Coxiella’ *lanes 8 - 10).* However, fractions in which mitochondrial protein was recovered were contaminated with additional organelles, demonstrated by markers for the lysosomes and the ER *(Figure 1B, ‘ER’ and ‘Lysosomes*’). Thus, this method was effective at separating *C. burnetii* and mitochondria, however, further purification was required to remove host cell contamination. To do this we employed magnetically conjugated monoclonal antibodies directed against human TOM22 (hTOM22), a receptor protein of the outer membrane TOM complex (*67*), to separate mitochondria from remaining cellular organelles.

**Figure 1:**
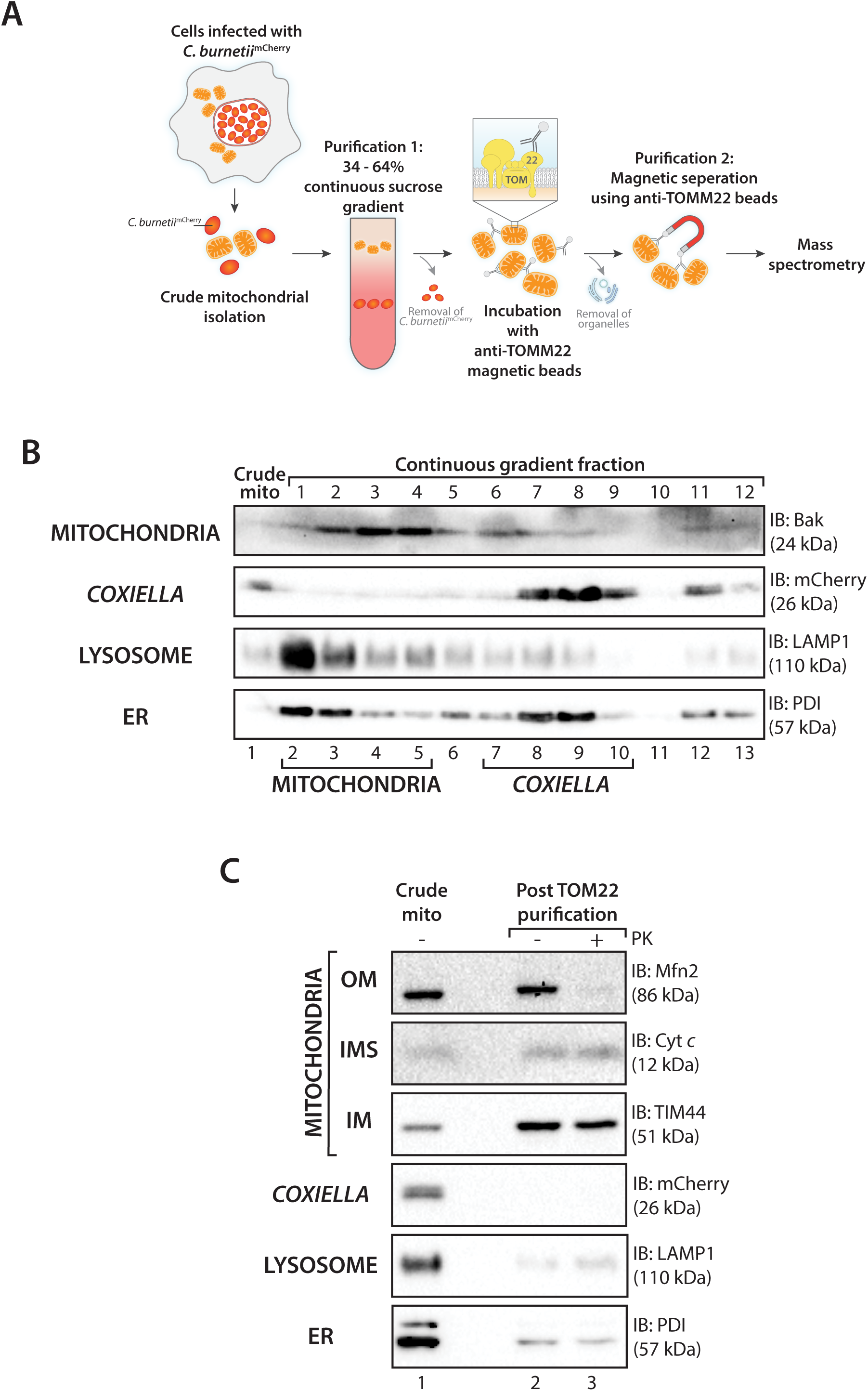
Mitochondrial purification from *C. burnetii-infected* cells. (A) Schematic representation of mitochondrial purification method from *C. burnetii-infected* cells. (B) Continuous sucrose gradient centrifugation of mitochondria isolated from *C. burnetii*_mCherry_-infected HeLa cells. A crude mitochondrial sample was taken pre-centrifugation (crude mito). Fractions were collected following ultracentrifugation and analysed by SDS-PAGE and immunoblotting with the indicated antibodies. Fractions containing the majority of mitochondrial or *C. burnetii* protein are marked with ‘mitochondria’ or *‘Coxiella’* respectively. (C) Mitochondria were isolated from THP-1 cells infected for 3-days with *C. burnetiimCheny* and purified by continuous sucrose gradient centrifugation and anti- TOMM22 separation. Samples were collected after crude mitochondrial isolation (Crude mito, *lane 1)* and after purification (Post TOMM22 purification, *lanes 2 and 3).* Purified mitochondria were incubated with or without Proteinase K (PK; 50 μg/mL) and analysed by SDS-PAGE and immunoblotting using the indicated antibodies as the listed organelle markers. OMM: outer mitochondrial membrane, IMS: intermembrane space, IMM: inner mitochondrial membrane, ER: endoplasmic reticulum, Cyt *c:* Cytochrome *c.*

Once the purification pipeline was established we wanted a system that would closely resemble the host cell environment during infection and therefore employed differentiated THP-1 cells for further analysis. THP-1 cells were infected with *C. burnetii* for 3 days and mitochondria isolated by differential centrifugation prior to purification by a continuous sucrose gradient followed by magnetic affinity purification. In comparison, mitochondria subjected to this dual purification approach (‘post TOM22 purification’) had an enrichment of mitochondrial proteins and a decrease in abundance of *C. burnetii*, lysosomes and the ER compared to the initial crude isolation (‘Crude mito’) (*Figure 1C, **lanes 1 and 2***). Addition of external protease, Proteinase K, to assess the integrity of the mitochondrial outer membrane resulted in loss of detectable signal for Mfn2, an outer membrane fusion mediator with a large cytosolic domain, while both cytochrome *c* (intermembrane space localised) and Tim44 (inner membrane localised) were protected from proteolytic degradation, indicating organelle integrity was maintained *(Figure 1C, lane 3).* The mitochondrial sample obtained following affinity purification had a reduction of *C. burnetii* as indicated by the lack of mCherry signal in the purified sample *(Figure 1C ‘Coxiella’).* Pure mitochondria were also less contaminated by lysosomes and the ER *(Figure 1C ‘lysosomes’ and ‘ER’).*

### Proteomic analysis of purified mitochondria from infected cells reveals depletion of *C. burnetii* proteins

Establishment of a method to isolate a pure mitochondrial fraction from infected cells provided an opportunity to address if *C. burnetii* effector proteins, in addition to MceA, localised to the organelle during infection. A time point of 3 days post-infection was chosen to capture an established stage of *C. burnetii* infection when the T4SS is known to be active (*19*). Mitochondria were isolated from *C. burnetii*-infected, differentiated THP-1 cells and purified using a continuous sucrose gradient and anti-hTOM22 affinity purifications *(Figure 1*). Equal amounts of sample were reserved at both the initial isolation and following the purification, designated ‘crude’ and ‘pure’ respectively *(Figure 2A*). Samples were analysed by label-free quantitative mass spectrometry and human and *C. burnetii* proteome coverage assessed *(Figure 2A*). Proteins were retained in the final dataset if they were identified in at least 2 of the 3 replicates analysed, the combined results of which identified >3000 proteins *(**Supplementary Table 1**).* Distinct proteome profiles were evident on comparing crude and pure replicates *(Figure 2B*). Encouragingly, proteins identified as bacterial clearly showed a reduction in abundance in the pure mitochondrial sample as compared to the crude (*Figure 2B, C. burnetii proteins highlighted in dark green*). Mitochondrial proteins were defined based on the human mitochondrial proteome database, MitoCarta *(Figure 2B, **Supplementary Table 1***, ‘*Mitochondrial’)* (*53, 54*). Quantitative comparison of the label free quantitation (LFQ) value of individual proteins within the crude and pure samples revealed the extent of the *C. burnetii* proteome coverage decreased ˜40% following purification confirming this method as valid in minimising *C. burnetii* present in the final mitochondrial sample *(Figure 2C*). Mitochondrial proteome coverage was mildly increased between the crude and pure samples (857 proteins listed in the human MitoCarta 2.0 mitochondrial database detected in crude, 900 detected in pure), indicating the purification technique was not detrimental to the organelle integrity *(Figure 2C*) (*53*). Gene ontology analysis for cellular compartment revealed proteins classed as neither mitochondrial nor bacterial and were identified in both the crude and pure samples. These proteins were annotated as residing in the ER, lysosomes or Golgi apparatus *(Figure 2C*). It is plausible that some of these proteins are from compartments in close association or contact with the mitochondrion such as the mitochondrial associated membrane (MAM). Importantly, these non-mitochondrial proteins were depleted in the pure sample compared to the crude, thus confirming the method as a robust approach to purify mitochondria from *C. burnetii-infected* cells.

**Figure 2:**
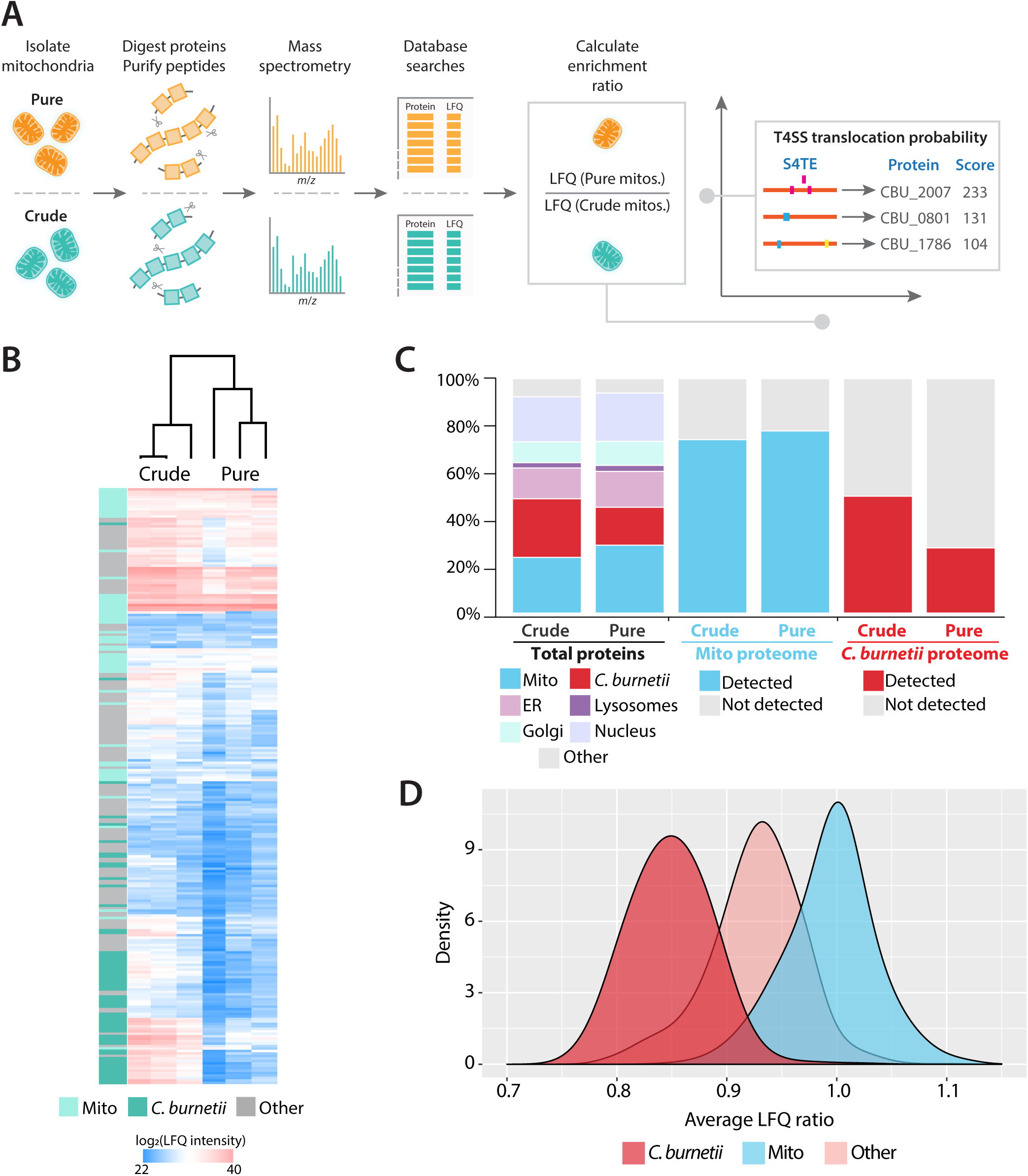
Proteomic analysis of mitochondria isolated from *C. burnetii-infected* THP-1 cells. (A) Overview of proteomic analysis pipeline. ‘Crude’ and ‘Pure’ mitochondrial samples were analysed by Label Free Quantitative (LFQ) mass spectrometry. LFQ values of individual proteins were compared to calculate the enrichment ratio. The enrichment ratio was then compared to the probability that a *C. burnetii* protein had of being an effector (T4SS score; https://sate.cirad.fr). (B) Heatmap and cluster analysis of ‘Crude’ and ‘Pure’ proteomics samples. Mitochondrial proteins are labelled in light teal, *C. burnetii* proteins are labelled in dark teal. n = 3 biological replicates per ‘crude’ and ‘pure’ fraction. Heatmap key denotes LFQ value (range 20 - 40). (C) Analysis of proteome coverage. Left columns: Total proteins identified across 3 biological replicates were classed as *C. burnetii* (Red; *C. burnetii* proteome), mitochondrial (Blue; MitoCarta 2.0 annotations) or endoplasmic reticulum (ER; pink), Golgi apparatus (Golgi; turquoise) or Lysosomal (Lysosomes; purple) according to Gene Ontology Cellular compartment annotation. Remaining proteins outside these categories were labelled ‘other’ (grey). Each number was compared to the total number of proteins identified in both the Crude and Pure mitochondrial preparations. Middle columns: Total mitochondrial proteins detected in Crude and Pure preparations were compared to the MitoCarta 2.0 mitochondrial proteome and coverage depicted. Right columns: Total *C. burnetii* proteins detected in Crude and Pure fractions were compared to the annotated *C. burnetii* proteome. (D) Density plot depicting the distribution of LFQ enrichment ratio (calculated as in (A)). Enrichment ratio of mitochondrial proteins (blue) and *C. burnetii* proteins (red) shown. Remaining proteins represented in pink.

### Analysis of protein enrichment reveals *C. burnetii* proteins associated with the mitochondria during infection

In order to determine the extent of protein enrichment as a result of purification, the LFQ values of host cell and bacterial proteins in the pure mitochondrial sample were compared to the crude, for each biological replicate *(Figure 2A, **Supplementary Table 1**: ‘LFQ pure/LFQ crude’).* This ratio was then averaged across biological replicates to yield a final ‘enrichment ratio’ (***Figure 2A***, ***Supplementary Table 1**: ‘LFQ ratio Average’*). This revealed higher LFQ ratios amongst mitochondrial proteins and therefore greater enrichment of mitochondrial proteins compared to *C. burnetii* proteins within the pure dataset (*Figure 2D, compare mitochondria [blue] to C. burnetii [red]*). These analyses indicate that our dual purification method enriches for mitochondrial proteins whilst decreasing the abundance of *C. burnetii* proteins within the pure sample.

Despite the rigorous method of purification and the reduction in co-purification of *C. burnetii* with mitochondria, bacterial proteins were still detected within the pure mitochondrial fraction (***Figure 2C***), showcasing both the sensitivity and challenges of mass spectrometry approaches. To distinguish between contaminating *C. burnetii* proteins and effector proteins we used the S4TE 2.0 Type 4 effector protein predictive tool (https://sate.cirad.fr) to generate a probability score of a *C. burnetii* protein being an effector (*57*). Analysis of the *C. burnetii* genome by the S4TE online program generated a list of 1818 proteins with an assigned score of translocation probability between 0 and 233 with a proposed effector score threshold >72 (***Supplementary Table 2***) (*57*). On comparison to the established cohort of 150 *C. burnetii* effector proteins, the S4TE output scores identified the majority of effector proteins although open reading frames unannotated in the National Centre for Biotechnology (NCBI) gene, nucleotide or protein database received either no score or a score below 72. The score for these proteins was manually annotated to 75 to account for this discrepancy *(**Supplementary Table 2**)*. To identify *C. burnetii* effector proteins associated with the mitochondria during infection, the enrichment factor was compared to the S4TE output (termed ‘T4SS probability’) *(Figure 2A, **Supplementary Table 2**)*. To ensure a high confidence list of proteins was produced, strict cut-offs for both parameters were applied: *C. burnetii* proteins were considered with an enrichment factor greater than 0.85, as this was above the enrichment factor mean of *C. burnetii* proteins identified and T4SS probability score greater than 72 to ensure a high-certainty of effector protein prediction. A cluster of 7 bacterial proteins were enriched alongside isolated mitochondria and had a high probability of being translocated by the T4SS (*Figure 3A, top right quadrant*). MceA (CBU0077), an effector protein previously identified and independently characterised as being targeted to the mitochondria, was identified alongside crude and purified mitochondria (***Supplemental Table 1***) (*28*). However, a low number of peptide sequences were associated with the protein in purified samples that did not culminate in valid LFQ values (***Supplemental Table 1***). This resulted in the effector protein removal from the final dataset. Nevertheless, identification of unique peptides from MceA in the purified mitochondria samples confirms this technique as a valid approach to identify candidate mitochondrial-targeted effector proteins. The 7 identified *C. burnetii* proteins exhibited a range of predicted biochemical properties, with diversity in size, predicted transmembrane domains (TMD), and the presence of signal peptide or mitochondrial targeting signals (*Figure 3B*). In addition to 5 effector proteins known to be translocated into the host cell (*24, 68, 69*) (*Figure 3A, proteins shown in green)*, our proteomics screen identified an additional 2 proteins that may represent candidate effector proteins *(Figure 3A, proteins shown in yellow).* Curiously, CBU1136 encodes multiple Sel-1-type TPR (SLR) motifs and annotated as enhanced entry protein C (EnhC) due to homology to the *L. pneumophila* protein of the same name (*70*). Although *C. burnetii* EnhC was found to contribute to bacterial virulence in a severe combined immunodeficiency (SCID) mouse model the function of this virulence factor in *C. burnetii* remains to be demonstrated (*71*). *L. pneumophila* EnhC inhibits soluble lytic peptidoglycan transglycosylase, suppressing NOD1-dependent NF-kB activation and is uniquely required for efficient intracellular replication (*72, 73*). Although, these 2 proteins represent an exciting direction of future research, for this study we focussed on the known effector proteins: CBU0937, CBU1425, CBU1594 and CBU1677 as CBU1863 has been previously characterised elsewhere (*69*).

**Figure 3:**
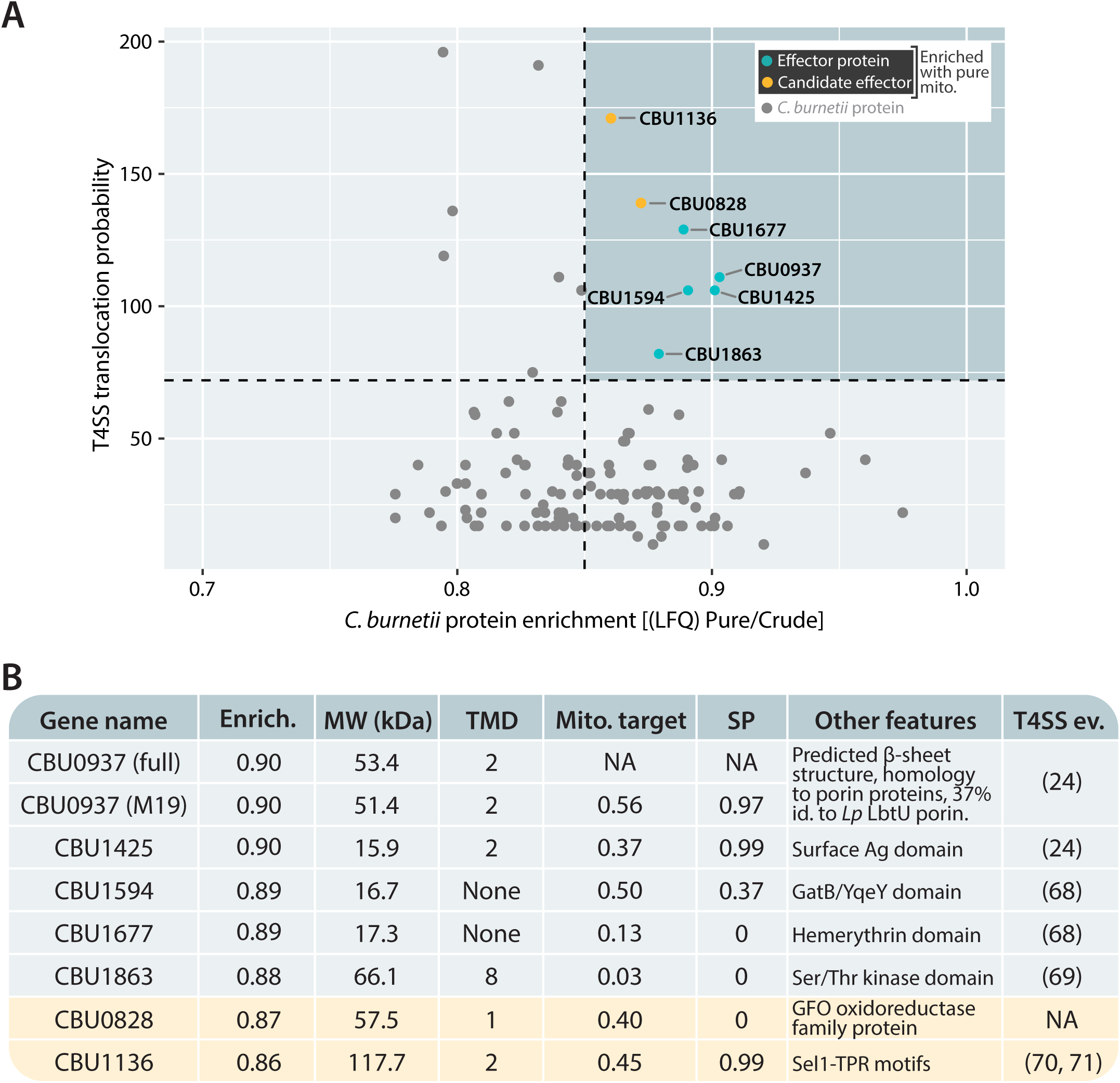
Analysis of *C. burnetii* proteins associated with purified mitochondria. (A) Scatter plot of *C. burnetii* protein enrichment factor (horizontal axis: LFQ (Pure/Crude)) compared to the probability of translocation by the T4SS (vertical axis). Dotted lines represent cut-offs: T4SS score > 72; enrichment factor > 0.85. Proteins with a high enrichment ratio and high probability of translocation shown in top right (blue quadrant). Proteins known to be translocated into the host cell are shown in green (‘effector protein’), proteins currently unannotated as effector proteins shown in yellow (‘candidate effector’). All other *C. burnetii* proteins depicted in grey. (B) Bioinformatic analysis of enriched *C. burnetii* proteins identified in top right quadrant of (A). ‘Enrich.’: enrichment ratio; TMD: Transmembrane domain; MW (kDa): molecular weight (kilodaltons); Mito. target: mitochondrial targeting signal prediction (determined using Mitoprot); SP: signal peptide prediction; Other features: additional features/homology to other proteins (determined using HHpred); Evidence/ref.: evidence/reference of T4SS-translocation or previously published studies regarding the protein.

### Confirmation of the subcellular localisation of mitochondrial-associated effector proteins

We set out to assess the validity of our results and complement our proteomic data with the ectopic expression and fluorescence microscopy of the identified effector proteins. This can provide valuable information into the subcellular localisation of effector protein in the host cell. Furthermore, ectopic expression of effector proteins supports downstream biochemical analysis of effector proteins, as often native effector proteins are expressed at very low levels in host cells. The selected effector proteins were epitope tagged (3XFLAG) at N or C termini and expressed in HeLa cells, since the monocyte-derived THP-1 cells exhibit an immune response to the introduction of plasmid DNA via transfection. We took into consideration that addition of the peptide tag may occlude targeting signals within the effector protein as well as the chance that additional effector proteins that are present during infection may act as chaperones to aid in effector targeting (*30, 74*). Therefore, the N and C-terminally tagged effectors were expressed in both uninfected and *C. burnetii-infected* HeLa cells (persistent infection). Transfections were performed over a shorter (18 hours) time period to capture protein localisation prior to gross over-expression. The host cell mitochondrial network was identified by staining for the matrix localised protein NDUFAF2, a Complex I assembly factor.

Expression of 3XFLAG fusion proteins in HeLa cells revealed a diversity in localisation patterns. The position of the 3XFLAG-tag influenced the localisation of the expressed proteins. Effector proteins with a 3XFLAG-tag located at the N-terminus of the protein, with the exception of CBU1425, did not demonstrate any clear co-localisation with the mitochondrial network *(Figure 4A*). A partial association with mitochondrial tubules was observed for _3XFLAG_CBU1425 *(Figure 4A*). The targeting of N-terminally tagged effector proteins was then assessed in the context of *C. burnetii* infection *(Figure 4B, CCV marked by* *). Consistently, _3XFLAG_CBU1425 localised in the vicinity of mitochondria *(Figure 4B*) while localisation differences were observed for _3XFLAG_CBU1677 *(Figure 4B*). _3XFLAG_CBU1677 expressed in uninfected cells was identified in vesicle-like structures located around the nucleus and condensed at opposing poles of the cell (***Figure 4A***). However, during *C. burnetii-* infection, transfected _3XFLAG_CBU1677 appeared to align with mitochondrial tubules in semi-regularly spaced foci *(Figure 4B*).

**Figure 4:**
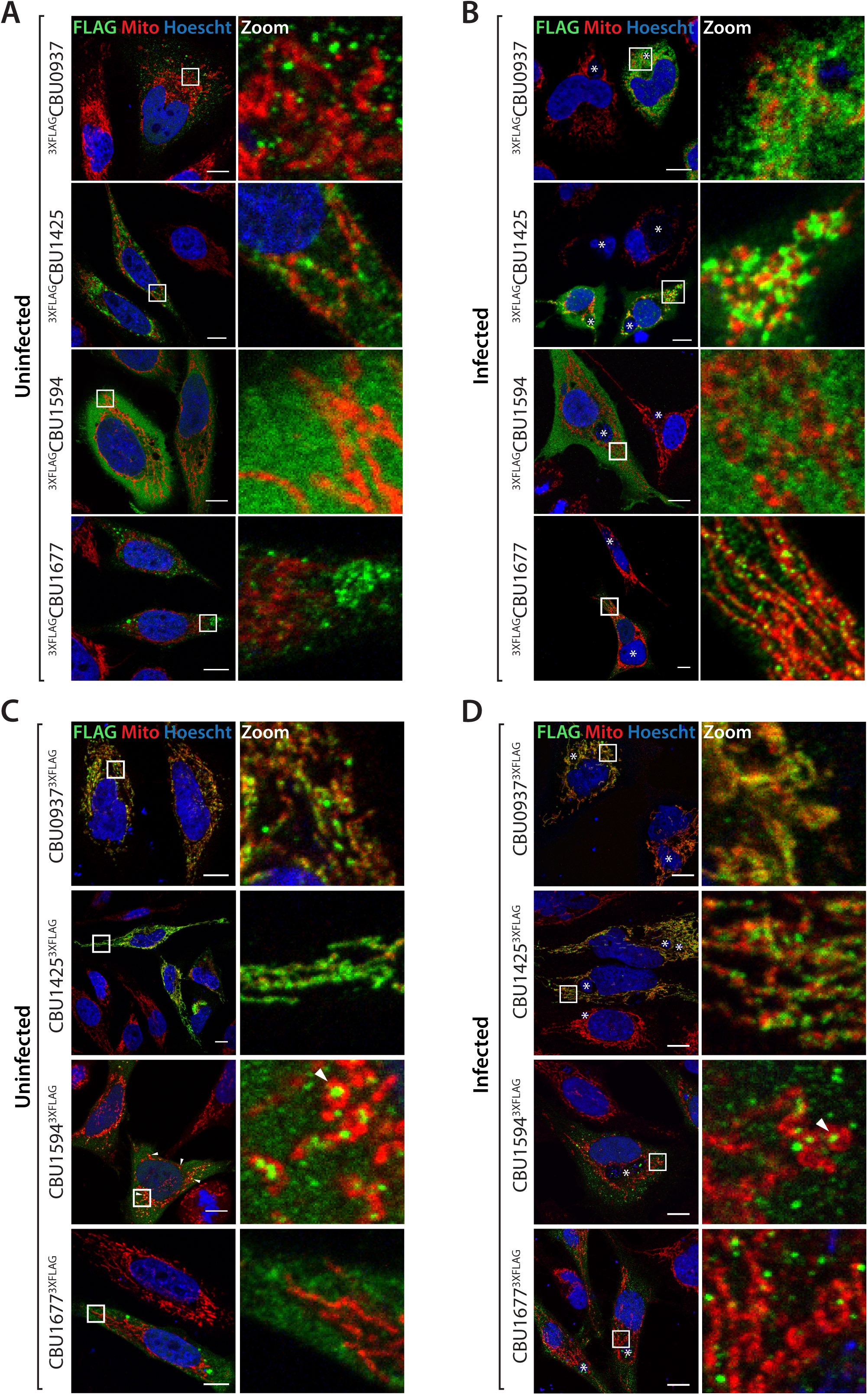
Localisation of N-terminally and C-terminally tagged, novel mitochondrial-targeted *C. burnetii* effector proteins. (A) Uninfected HeLa cells were transfected for 18 hours with N- terminally 3XFLAG-tagged effector protein. (B) *C. burnetii*-infected HeLa cells were transfected for 18 hours with N-terminally 3XFLAG-tagged effector protein. (C) Uninfected HeLa cells were transfected for 18 hours with C-terminally 3XFLAG-tagged effector protein. (D) Infected HeLa cells were transfected for 18 hours with C-terminally 3XFLAG-tagged effector protein. All cells were fixed and immunostained with antibodies against FLAG (green) and NDUFAF2 (Mito; red). Nucleus was stained with Hoechst 33258 (blue). Cells were imaged by confocal microscopy. Scale bar represents 10 μm. Right panels show a magnified view of the boxed region (‘zoom’). * denotes position of CCV. Arrowheads denote regions of interest.

Next, the localisation of effector proteins tagged at the C-terminus was established. The altered position of the 3XFLAG tag had a profound effect on the observed localisation of CBU0937, CBU1425 and CBU1594 *(Figure 4C*). In contrast to the absence of any distinct localisation of _3XFLAG_CBU0937, CBU09 37_3XFLAG_ co-localised with the host cell mitochondrial network, distributed along mitochondrial tubules *(Figure 4C*). Similarly, CBU14253XFLAG was also found to co-localise with the signal from the mitochondrial marker NDUFAF2 *(Figure 4C*). CBU15943XFLAG localised to distinct, punctate structures that were in the vicinity of mitochondria (***Figure 4C***). Intriguingly, these CBU1594_3XFLAG_ puncta were often found in the middle of mitochondrial tubules that had curved into donut-shaped structures *(Figure 4C, white triangles).* CBU16773XFLAG demonstrated a similar localisation pattern as the N-terminally tagged equivalent (_3XFLAG_CBU1677) in uninfected cells, with concentrations of protein near to the nucleus *(Figure 4C*).

Localisation of C-terminally tagged effector proteins in *C. burnetii-infected* cells revealed, as for uninfected cells, CBU0937_3XFLAG_ and CBU1425_3XFLAG_ co-localised with the mitochondrial network *(Figure 4D, CCV marked by* *). CBU1594_3XFLAG_ was again located in punctate structures close to mitochondria, occasionally encircled within mitochondrial tubules (*Figure 4D, white triangles*). Similar to the N-terminally tagged protein expressed in infected cells, CBU16773XFLAG localised to vesicular structures in close association with mitochondria, indicating that the targeting information for this protein possibly requires additional factors (presumably other effector proteins) only present during infection *(Figure 4D*).

Taken together, the findings of our proteomic screen identified 7 novel proteins associated with the mitochondrion during infection and microscopic analysis of 4 of these proteins supports their mitochondrial localisation: CBU0937, CBU1425, CBU1594 and CBU1677. We therefore propose to rename these proteins mitochondrial *Coxiella* effector proteins B (CBU0937), C (CBU1425), D (CBU1594) and E (CBU1677). These proteins display some variation in localisation at the mitochondria, from a more uniform distribution along the mitochondrial tubules as for MceB and MceC to more punctate structures such as MceD and MceE *(Figure 4*). Variation in targeting of these proteins was observed dependent on the location of the tag and the context of infection, providing valuable insight into the targeting information required for protein localisation.

### Characterising the role of MceC - a C. *burnetii* effector protein targeted to the mitochondria during infection

Given the clear localisation of MceC with the mitochondrial network we decided to investigate this protein further. CBU1425 is a 15.9 kDa protein, predicted to contain a signal peptide and 2 TMDs, located towards the N-terminus *(Figure 5A*). Sequence analysis of MceC revealed the protein contains multiple glycine zipper (GXXXG or GGXXG) motifs spanning residues 27 - 69 and encompassing the 2 TMDs *(Figure 5A*). Glycine-zipper motifs are often found in TMDs and typically assist in mediating protein-protein interactions such as homo-oligomerisation and helix packing within a membrane (*75*). The C-terminal part of the protein contains a region with 42% homology to a ‘17 kDa outer membrane surface antigen’ domain present in some Proteobacterial species *(Figure 5A*). BLAST searches do not reveal significant homology to any mammalian proteins, which is common for *C. burnetii* effector proteins. Earlier studies focused on uncovering *C. burnetii* T4SS substrates demonstrated MceC was delivered into the host cell by the T4SS using a BlaM reporter assay (*24*). The subcellular localisation of the protein has been previously suggested to be mitochondrial, however, no experimental evidence for this phenotype was provided (*68*).

**Figure 5:**
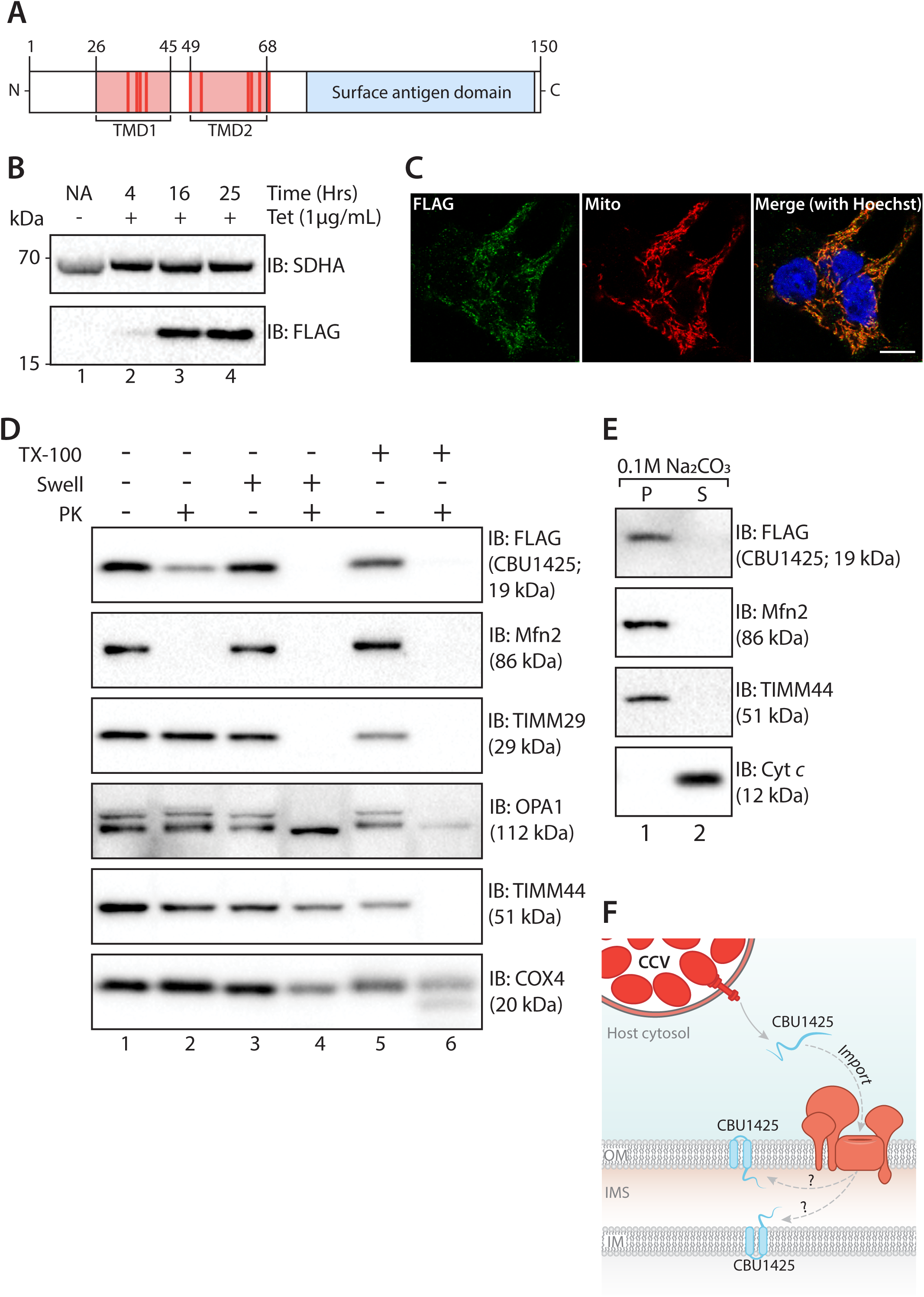
Characterising the submitochondrial localisation of MceC. (A) Schematic representation of predicted domain structure of MceC. Location of predicted transmembrane domains (pink), glycine motifs (red) and region of homology to ‘surface antigen domain’-containing proteins (blue) depicted. Numbers denote amino acid locations. (B) Protein expression in tetracycline inducible HEK293MceC-3XFLAG cells. Cells were left untreated (NA) or treated with tetracycline (1 μg/mL) for 4, 16 or 25 hours to induce expression of MceC_3XFLAG_. Cells were collected and analysed by SDS-PAGE and immunoblotting with the indicated antibodies. (C) HEK293MceC-3XFLAG cells were treated for 16 hours with tetracycline (1 μg/mL) before being fixed and stained with antibodies against FLAG (green) and NDUFAF2 (Mito; red). Nucleus was stained with Hoechst 33258 (blue). Scale bar represents 10 μm. (D) Mitochondria were isolated from HEK293 cells expressing MceC_3XFLAG_ and either left intact, (lanes 1 and 2), subjected to hypotonic swelling (‘Swell’) of the outer mitochondrial membrane (mitoplasts; lanes 3 and 4) or solubilised mitochondria (TX-100; lanes 5 and 6). Samples were left untreated or incubated with Proteinase K (PK; 50 μg/mL, lanes 2, 4 and 6) before analysis by SDS-PAGE and immunoblotting with the indicated antibodies. (E) Mitochondria isolated from HEK293 cells expressing MceC_3XFLAG_ were subjected to alkaline extraction in sodium carbonate (Na2CO3; 0.1M, pH 11). Following incubation on ice, membrane (P) and soluble (S) fractions were separated by ultracentrifugation prior to SDS-PAGE and immunoblotting with the indicated antibodies. (F) Proposed sub-mitochondrial localisation of MceC.

A stable tetracycline-inducible HEK293T cell line expressing MceC_3XFLAG_ was created, protein expression induced for 4, 16 and 25 hours and expression analysed by immunoblotting *(Figure 5B*). As early as 4 hours MceC_3XFLAG_ could be detected and a steady increase in expression was evident, whilst little to no change was apparent in the levels of the loading control, the Complex II subunit SDHA, located in the inner membrane *(Figure 5B*). At 16 hours induction, MceC_3XFLAG_ was observed to co-localise with the mitochondrial marker NDUFAF2, confirming the expression of the protein detected by immunoblotting and correct protein targeting *(Figure 5C*). As the levels of bacterial effector proteins are typically very low within cells it was important to determine that the level of MceC expression was not causing protein misfolding and/or aggregation. We conducted solubility trials across a range of different non-ionic detergents 4 and 8 hours post protein induction with tetracycline. Isolated mitochondria were resuspended in digitonin (1% (w/v)), TX-100 (1% (v/v)) or DDM (0.2% and 0.4% (w/v)) and analysis of a total, pellet and supernatant fraction by SDS-PAGE and immunoblotting revealed that MceC_3XFLAG_ was recovered in the supernatant fraction following 4 hours of induction *(**Supplemental Figure 1A, lanes 3, 6, 9 and 12**).* Samples induced for 8 hours displayed the protein in the pellet during the digitonin condition, indicating the protein was insoluble *(**Supplemental Figure 1B**)*, therefore, protein expression for 4 hours was selected for further experiments.

Firstly, we assessed which mitochondrial sub-compartment MceC_3XFLAG_ was localised within by performing a mitochondrial subfractionation assay. Mitochondria isolated from HEK293 expressing MceC_3XFLAG_ cells, were left untreated, subjected to swelling in hypo-osmotic buffer or completely solubilised in TX-100 *(Figure 5D*). The accessibility of proteins in each of these conditions to Proteinase K was then assessed prior to analysis by SDS-PAGE and immuno-decoration for proteins of distinct mitochondrial sub-compartments *(Figure 5D*). The FLAG signal, corresponding to MceC_3XFLAG_, was inaccessible to Proteinase K in intact mitochondria, however, protection was lost following the rupture of the OMM during swelling *(Figure 5 D, compare lanes 2 and* 4). Upon comparison, this pattern of degradation is similar to the inner membrane protein Tim29, a component of the TIM22 complex, but distinct from the inner membrane/matrix Complex IV subunit, COX4 *(Figure 5D*). These results indicate that the C-terminus (location of 3XFLAG-tag) of MceC_3XFLAG_ is located within the intermembrane space of the mitochondria. We also noted a drop in the intensity of FLAG signal upon treatment of intact mitochondria with Proteinase K *(Figure 5D, compare lanes 1 and* 2). This could be attributed to the collection of pre-imported MceC_3XFLAG_ at the outer membrane, as the expression level in the inducible cell system, whilst regulated, is still likely higher than the endogenous levels of protein during infection.

As MceC is predicted to contain 2 TMDs, the integration of the protein into a mitochondrial membrane was assessed using carbonate extraction. We reasoned that MceC could be located within either the outer or inner membrane whilst still maintaining the C-terminus in the intermembrane space and thus the 3XFLAG-tag accessible for degradation *(Figure 5D*). Mitochondria isolated from HEK293 cells expressing MceC_3XFLAG_ and were resuspended in sodium carbonate for separation of membrane proteins and peripheral membrane proteins by high-speed centrifugation *(Figure 5E*). MceC_3XFLAG_ was localised to the pellet fraction, alongside Mfn2 and Tim44, indicating that the protein is indeed integrated into a mitochondrial membrane with the C-terminus in the intermembrane space *(Figure 5F*).

### MceC can interact with components of the mitochondrial quality control machinery

Effector proteins typically act to modulate a specific host process. This often involves a direct interaction with host cell proteins. We sought to identify any mitochondrial interacting partners of MceC by immunoprecipitation of MceC_3XFLAG_. MceC_3XFLAG_ and associated proteins were purified by FLAG antibodies and eluates separated on SDS-PAGE to confirm efficiency of pull-down (***Figure 6A***). A signal corresponding to MceC_3XFLAG_ was detected using FLAG antibodies in the elution fraction of HEK293MceC-3XFLAG cells but not control cells *(Figure 6A, lanes 5 and 10).* Accordingly, the mitochondrial protein SDHA was not detected in the elution *(Figure 6A, lane 10).* When eluates from 3 independent, biological replicates were processed for label-free quantitative mass spectrometry we observed a successful enrichment of MceC_3XFLAG_ in addition to several other mitochondrial proteins *(Figure 6B, C and **Supplemental Table 3**)*. Intriguingly, these proteins included important players in the mitochondrial proteostasis and quality control pathways YME1L, SCO2, CLPB and SLP2 as well as components of the TOM complex: TOM20 and TOM40 (*Figure 6B, **C** and **Supplemental Table 3***). Additionally, key proteins involved in inner membrane and cristae morphology, OPA1 and the MICOS subunit MIC19 were also enriched although not above the significance threshold *(Figure 6B, **C** and **Supplemental Table 3**)*.

**Figure 6:**
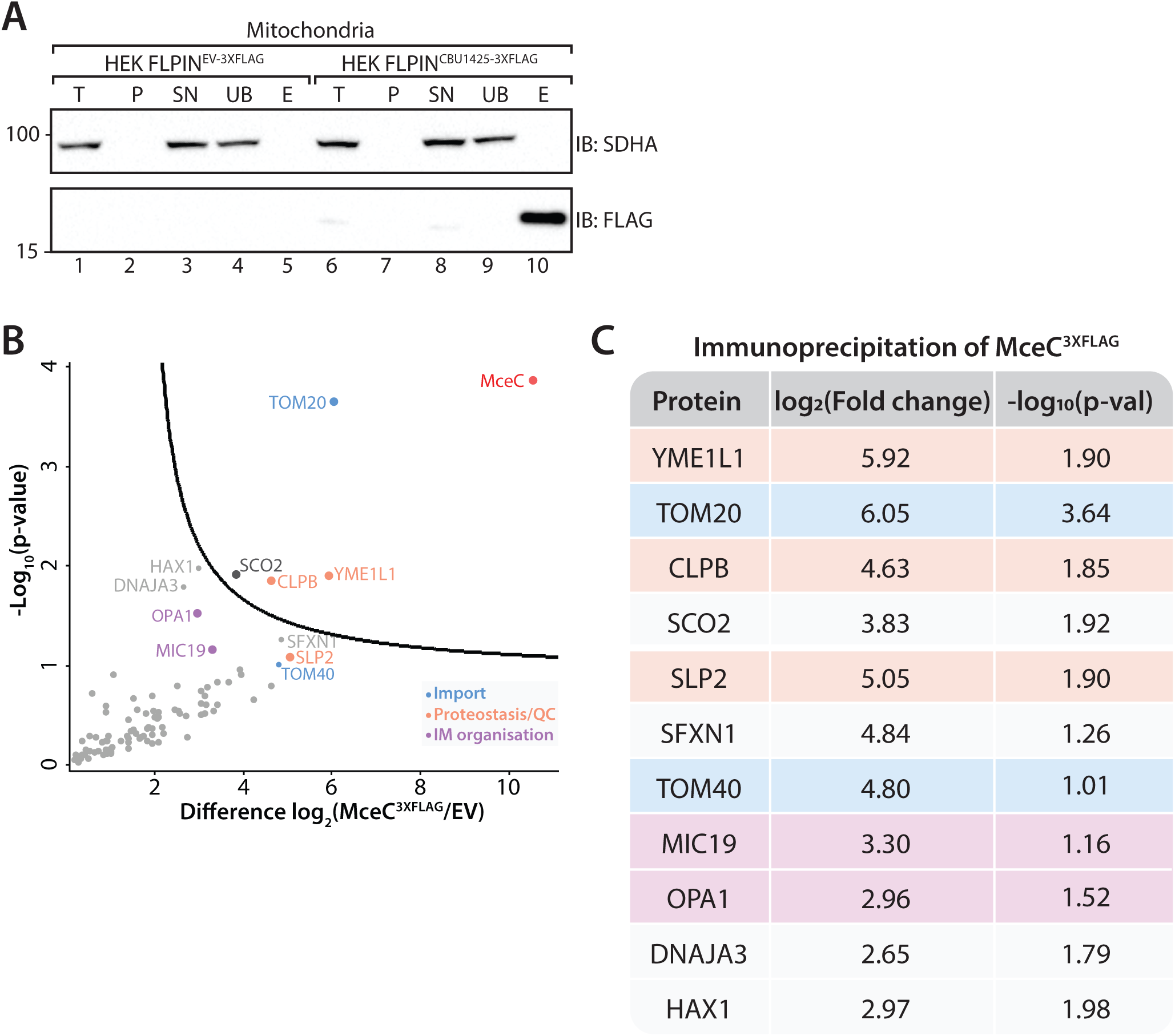
MceC immunoprecipitates resident mitochondrial proteins. (A) Mitochondria isolated from control and HEK293 cells expressing MceC_3XFLAG_ for 4-hours were solubilised in digitonin (1% (w/v))-containing buffer and lysates subjected to anti-FLAG immunoprecipitation. Collected fractions were analysed by SDS-PAGE and immunoblotting with the indicated antibodies. T: Total, P: Pellet, SN: Supernatant, UB: Unbound, E: Elution. 10% of the T, P, S and UB fractions and 100% of the elution fraction was loaded for analysis. (B) Volcano plot showing proteins enriched following MceC immunoprecipitation as outlined in (A) compared to control ‘empty vector’ (EV) cells. Mitochondrial proteins plotted and each circle represents one protein. Gene names of selected proteins are used for labels. Horizontal axis shows the Log_2_(fold change) of MceC interacting partners and vertical axis shows -Log_10_(p-value) of two-sample Students T-test (FDR: 0.05, s0: 1). n = 3 biological replicates. QC: Quality Control, IM: inner membrane. Colours correspond to key. (C) Proteins identified by quantitative mass spectrometry following immunoprecipitation of MceC in (B). Gene names of selected proteins used as protein label. Corresponding Log_2_(fold change) and Log_10_(p-value) of each protein listed. Colours as per key in (B).

To assess if any of these interactions were conserved during infection, HEK293_MceC-3XFLAG_ and control cells were persistently infected with *C. burnetii* prior to protein induction. Mitochondria were isolated, solubilised in digitonin and immunoprecipitation with FLAG antibodies performed before analysis by mass spectrometry. Consistent with the interactions observed in uninfected cells, SLP2 was enriched alongside MceC_3XFLAG_ in mitochondria isolated from infected cells (***Supplemental Figure 2 and Supplemental Table 4***). Interaction with YME1L was not observed in infected cells and may indicate CBU1425 interaction with the protease is less stable during infection.

### Impact of MceC expression on mitochondrial function

To gain perspective on the wider implications of MceC localisation to the mitochondria we looked more broadly at changes to the mitochondrial proteome over the expression time-course. To investigate this, label free quantitative mass spectrometry was performed on mitochondria isolated following 1, 4 and 8 hours MceC_3XFLAG_ expression. Triplicate samples were compared using an analysis of variance (ANOVA) test to determine significantly altered protein abundance at each time point. This revealed 93 proteins with an FDR of 5% and S0 of 0 *(Figure 7A, **Supplemental Table 5***). These 93 proteins were involved in a range of mitochondrial processes including metabolic functions including 3 solute carrier proteins of the inner mitochondrial membrane (glutamate carrier 1, the calcium-dependent aspartate-glutamate carrier and the ATP-Mg/Pi carrier), translation and import including Tim29, the mitochondrial intermediate peptidase MIPEP and inner membrane translocase TIM23 motor associated subunit Pam16. Considering the probable inner membrane localisation of MceC, the effector is in an advantageous position to modulate many of these processes. These observations using a regulated protein expression system provide a unique snapshot of the impact of MceC localisation to the mitochondria in the absence of the full assault of *C. burnetii*-infection and additional effector proteins.

**Figure 7:**
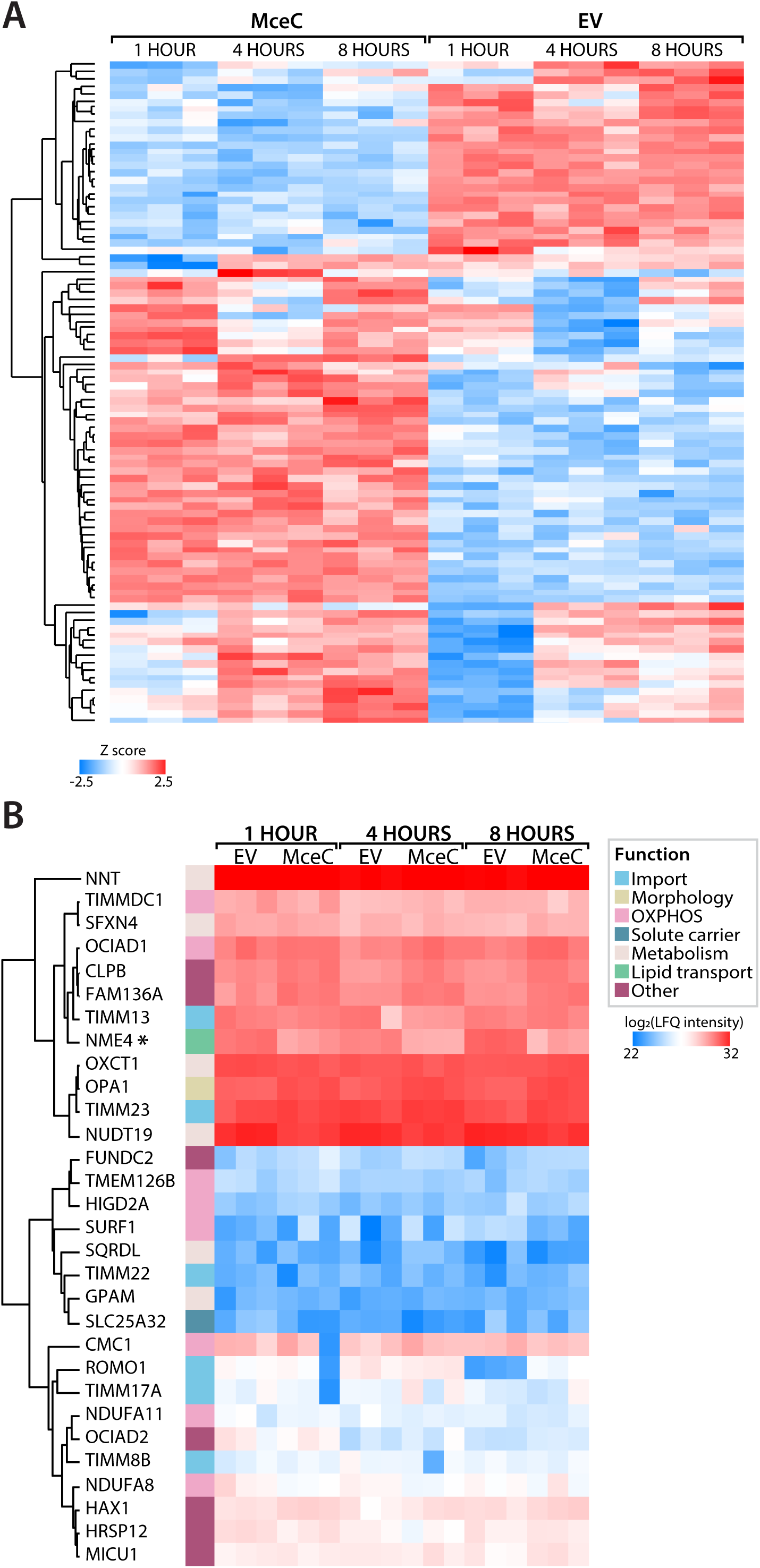
Expression of MceC affects mitochondrial function. Quantitative proteomics analysis of mitochondria isolated from control (EV) or MceC-expressing cells following 1,4 or 8 hours (HRS) tetracycline-induction. n = 3 biological replicates. (A) Heatmap of Z-scored log_2_-transformed label free quantitation (LFQ) intensities of proteins identified as significantly altered across all time points. (B) Heatmap of log_2_-transformed label free quantitation (LFQ) intensities of known YME1L substrate proteins. Gene names of selected proteins used as labels. Proteins labelled according to function and colours correspond to key.

Of the interacting partners enriched within MceC immunoprecipitates, YME1L was particularly interesting. YME1L forms a hexameric, ATP-dependent proteolytic complex in the inner membrane that has emerged as a central regulator of mitochondrial biogenesis, coupling protein quality control with organelle dynamics and function (*76*). Given the diverse and important roles of YME1L within mitochondria we decided to further investigate the impact of MceC on YME1L function. When LFQ values of known substrates of YME1L where compared, the vast majority of YME1L substrates underwent little to very subtle changes over the time examined *(Figure 7B, Supplemental Table* 5). Curiously, only one YME1L substrate, the inner membrane localised nucleoside diphosphate kinase NME4, was significantly altered with a decreasing trend across all time points *(Figure 7B, denoted by* *). Over this analysis, NME4 had a fold change ratio of −1.61 (p-value: 0.004) following 1 hour MceC expression, which decreased to a fold change of −2.17 (p-value: 0.002) at 8 hours *(Supplemental Table 5)* Grouping of these substrate proteins according to their mitochondrial function did not reveal any collective changes to a mitochondrial process, however, it is interesting to note that in addition to being significantly altered across all time-points, NME4 was the only protein attributed to lipid transport *(Figure 7B*). Thus, the interaction of MceC with YME1L does not appear to lead to a broad change in the proteolytic capacity of YME1L and may instead be a more targeted regulation of a subset of proteins important to *C. burnetii* infection.

## Discussion

The development of techniques to endogenously study host-pathogen interactions is integral for the progression of the field. Adaptation of high-sensitivity technology such as super resolution microscopy, mass-spectrometry and electron cryotomography to detect and determine structure of a protein or protein complexes, present exciting opportunities to advance the toolkit available to understand how pathogens manipulate the host cell during infection. In this study we develop a proteomics-based screening approach to identify *C. burnetii* effector proteins targeted to distinct sub-cellular compartments and show the application of this technique to identify mitochondrially-targeted effector proteins. Mass-spectrometry presented an elegant tool to achieve this as the development of high-sensitivity mass-spectrometers now allows identification of proteins that exist in relatively low abundance, such as an effector protein amidst the host cell proteome (*77*). This is the first study to utilise a broader, unbiased and unlabelled approach to capture the interactions of native *C. burnetii* proteins with the host cell mitochondrion. Development of these methodologies is particularly important for pathogens with large effector repertoires where a key outstanding question remains: are these proteins *bona fide* effectors that are translocated into the host cell during infection?

This non-targeted approach allowed the identification of 7 proteins associated with the mitochondria during natural infection. Of the 7 *C. burnetii* proteins associated with the organelle during infection, 5 have already been shown to be translocated by the T4SS into the host cell (*24, 78*). Whilst we focused on the known effector proteins, the high enrichment of CBU0828 and CBU1136 (EnhC) suggests these proteins may represent important virulence factors during intracellular *C. burnetii* infection (*71-73*). As the established cohort of *C. burnetii* effector proteins is based on over-expression reporter systems, this approach also validates the native proteins are indeed effectors. Of the known effector proteins further investigated for mitochondrial localisation, CBU0937, CBU1425, CBU1594 and CBU1677 displayed association with the mitochondria validating their proteomic enrichment with the organelle. Curiously, transposon mutagenesis screening has demonstrated that CBU0937 makes a significant contribution to the intracellular replication of *C. burnetii* further validating the importance of the host cell mitochondrion during infection (*68*). Further characterisation of CBU1425 revealed that the protein is imported and integrated into a mitochondrial membrane, likely the inner membrane and may influence organelle quality control systems. Thus, we propose the renaming of these 4 effector proteins to Mitochondrial *Coxiella* effector protein B - E (MceB - E). These proteins join the previously identified mitochondrial-localised effector MceA, forming a cohort of *C. burnetii* proteins that target the organelle during infection.

Our investigation focused on the interaction between *C. burnetii* and the host cell mitochondrion, developing a reliable and robust method to separate the organelle from both bacterial and host cell contamination. We propose the techniques and analytical approaches developed here could be applied to the study of other organelles during *C. burnetii* infection, or more broadly, to the study of other intracellular bacterial pathogens. Whilst previous studies have been undertaken to look at bacterial containing vacuoles, few have attempted to look at organelles free of bacterial contamination during infection (*79, 80*). Expanding this study spatially and temporally would provide a systematic, comprehensive analysis of the effector protein localisation and dynamics within the host cell over the full course of *C. burnetii* infection. Studies performed on viral pathogens such as human cytomegalovirus, have provided an integrated insight into host organelle composition as well as viral protein targeting and complex formation during infection (*81, 82*). This also indicated viral protein translocation between cellular compartments during infection (*81, 83*). If, or when bacterial effector proteins traffic between multiple organelles during infection is not well understood. Investigations into the function of *C. burnetii* effector AnkG, demonstrated the protein is capable of trafficking between the mitochondria and the nucleus via a ‘piggy-backing’ mechanism to interrupt nuclear apoptotic signalling (*38, 84*). Whilst our proteomics analysis did not capture the enrichment of AnkG at the mitochondrion, it is possible that this interorganelle trafficking had already resulted in translocation of the protein to the nucleus. Considering *C. burnetii* infection persists in the host cell for approximately 7 days, it is possible that effector proteins could be dually localised or localised to membrane contact sites between organelles at distinct stages of infection. Our increasing understanding surrounding the interaction outcome of effectors of effectors (‘meta-effector’) has revealed how effector proteins may be regulated post-translationally (*11*). It will be interesting to see if this modulation may also affect effector protein targeting within the host cell. Therefore, spatial and temporal application of the techniques developed here could form a ‘road map’ of effector targeting and trafficking events that could assist in functionally distinguishing between effector proteins, a particular advantage in bacterial species with large effector protein repertoires such as *C. burnetii.*

Purification of the mitochondrion by the method presented here, whilst effective, requires the removal of the organelle from the host cell. This and downstream processing could disrupt transient effector-host protein at the organelle surface or at inter-organelle contact sites. The use of *in situ* protein proximity labelling approaches such as APEX has considerably broadened our understanding of the mitochondrial proteome (*85, 86*). Application of this technique during the study of intracellular pathogens could provide insight into both the spatial and interaction characteristics of effector proteins localised to the mitochondria or associated membranes. Recently, improvement of an affinity-enrichable, isotopically coded, mass spectrometry-cleavable crosslinking approach coupled to a targeted MS2 acquisition strategy and data analysis allowed the identification of unique protein-protein interactions within the yeast mitochondrial proteome (*87*). The ability to cross-link effector-host protein interactions *in-organello* could be particularly advantageous to the study of effector interactions without the need for overexpression systems. Conversely, incorporation of a non-canonical amino acid into bacterial proteins allows the specific labelling of bacterial proteins, distinguishable from the host cell proteome (*88-90*). The click-chemistry based enrichment of bacterial proteins during infection could greatly assist in identifying effector proteins in low abundance during intracellular infection.

Introduction of bacterial proteins into the host cell by the T4SS can impact how an effector protein behaves within the cell. Structural characterisation of the *L. pneumophila* T4SS has provided valuable insights into deciphering the translocation mechanism underlying this complex, multi-protein apparatus (*8, 9*). What is clear from these structural studies is that delivery of a native, T4SS- translocated effector protein is distinct to protein targeting through the ER or secretory pathway as may occurs during ectopic expression. This was evidenced previously during our characterisation of MceA (*28*). With acknowledgement of these central caveats, proteins of interest were examined with both N and C terminal tags in both uninfected and infected cells. Indeed, this comparison revealed interesting aspects of effector protein biology. For instance, MceE (CBU1677) demonstrated distinct localisation only when expressed in infected cells, indicating a potential role for additional *C. burnetii* proteins in aiding this targeting. Furthermore, mitochondrial targeting of MceB (CBU0937) and MceC was observed only when the N terminus of the protein was exposed, suggesting the mitochondrial targeting information for these effector proteins is located in this region. Bacterial effector proteins frequently hijack host targeting signals to direct the protein to specific cellular locations (*32, 91*). In the case of the mitochondrion, the evolutionary relationship between the organelle and Gram-negative bacteria could be exploited to aid in effector protein targeting. Although mitochondrial protein import is a highly specialised process, several lines of evidence indicate that bacterial proteins are capable of utilising the mitochondrial import machinery for entry to and sorting within the organelle (*92-98*). Bioinformatic analysis of the domain structure of MceB predicts the protein to contain p-sheet topology and homology to bacterial porin proteins. β-barrel proteins of *Neisseria sp.* PorB and Omp85 have been shown to successfully utilise the mitochondrial import machinery for import and assembly into mitochondrial membranes (*95, 96, 99*). PorB association with the host cell mitochondria disrupts the membrane potential, sensitising cells to apoptosis (*95*). Sequence analysis indicates that MceB lacks the classical β-targeting signal of mitochondrial β-barrel proteins (*100*). Considering the homology it has to porin-like proteins, it will be interesting to determine the precise interaction of this effector protein with human mitochondria during C. *burnetii* infection.

Prior to this study, MceC was a functionally uncharacterised *C. burnetii* effector protein (*24*). Mitochondrial subfractionation and carbonate extraction revealed the protein is imported into the organelle and integrated into a mitochondrial membrane, with the C-terminus of the protein in the IMS. MceC does not contain a classical mitochondrial targeting sequence (MTS), nor does it contain a mitochondrial intermembrane space targeting signal, raising the question how the effector protein reaches this location (*32*). Curiously, the substrate spectrum of the human translocase of the inner mitochondrial membrane (hTIM22) complex was recently expanded and the complex was identified as capable of importing non-canonical substrates, containing 2 or 3 transmembrane segments (*101*). It is interesting to postulate that *C. burnetii* may exploit a non-canonical route to import MceC into mitochondria. Immunoprecipitation of MceC identified several, resident mitochondrial interacting partners including several components of the mitochondrial quality control machinery, namely the *i-AAA* protease, YME1L and the caseinolytic peptidase, CLPB. Proteins of the mitochondrial quality control network include chaperones and proteolytic enzymes that survey and maintain mitochondrial homeostasis in response to organelle and cellular stresses (*102, 103*). Although in our system, MceC was expressed at a low level, we cannot rule out that these interactors are responding to the presence of MceC and attempting to remove the protein to maintain organelle proteostasis. However, MceC also immunoprecipitates additional components of the YME1L quality control network including SLP2, a component of the SLP2-YME1L-PARL (SPY) proteolytic complex and OPA1 (*104*). Additionally, global changes to the mitochondrial chaperone and proteostasis pathway were not evident over the 8 hour MceC expression proteomics time course. Notably, immunoprecipitation of MceC in the context of *C. burnetii* infection revealed conservation of the interaction with SLP2 but not YME1L. These observations advocate away from a scenario in which interaction of MceC with these mitochondrial proteins represents a quality control response, however this needs to be experimentally verified.

The initial characterisation of the mitochondrial targeting and interactions of MceC raises the question as to what role this effector protein has during *C. burnetii* infection. Recently, additional YME1L substrates were identified using quantitative proteomics in *Ymell*_−/−_ MEF cells (*105*). Studying the impact of MceC on the mitochondrial proteome can hint more broadly at the role of the effector protein in the cell, further extending the use of proteomics in the functional study of effector proteins during infection. Using proteomics analysis coupled to a time-course of MceC expression, we were able to rule out the collective modulation of a functional subset of YME1L substrates. Regulation of mitochondrial dynamics due to processing of the dynamin-like GTPase OPA1 is perhaps one of the best characterised roles of YME1L (*76, 106*). Changes to OPA1 processing has been implicated in altering mitochondrial morphology in response to different cellular stresses, metabolic and environmental demands (*105, 107, 108*). Whilst mitochondrial morphology has been reported to be modified during infection with *Listeria monocytogenes, Vibrio cholerae* and *L. pneumophila*, thus far, no study has demonstrated any mitochondrial network alterations during *C. burnetii* infection (*35, 109-111*). Although our analysis revealed no change to level of OPA1 over 8 hours of MceC expression, we cannot exclude changes in the abundance of the proteolytically processed OPA1 variants (*76*). Investigation into the abundance of these variants and of the mitochondrial inner membrane ultrastructure by electron microscopy during MceC expression and more broadly *C. burnetii*-infection could reveal a more complex role for MceC within the network of inner membrane quality control. Understanding the importance of MceB - E to *C. burnetii* infection by the characterisation of *C. burnetii* mutants lacking these genes could provide further clues to the proteins function during infection and in response to various cellular stresses. Determining any alterations to the mitochondrial proteome during infection in the absence of the effector protein with a *C. burnetii* strain lacking *cbu1425* could be particularly insightful given the interaction profile identified here. Conversely, analysis of *C. burnetii* replication in human cell lines lacking MceC interacting partners could also shed light on the significance of mitochondrial processes during *C. burnetii* infection.

This research has showcased the power of an unbiased proteomics-based technique to investigate and understand the host-pathogen interactions occurring between the host cell mitochondrion and the intracellular bacterial pathogen *C. burnetii.* This has established a method to investigate effector protein targeting during native infection and revealed 4 additional, mitochondrially-targeted *C. burnetii* effector proteins. The techniques developed here demonstrate the feasibility of applying mass spectrometry to create a map of the sub-cellular dynamics of native effector proteins during intracellular bacterial infections.

**Supplemental Figure 1:**
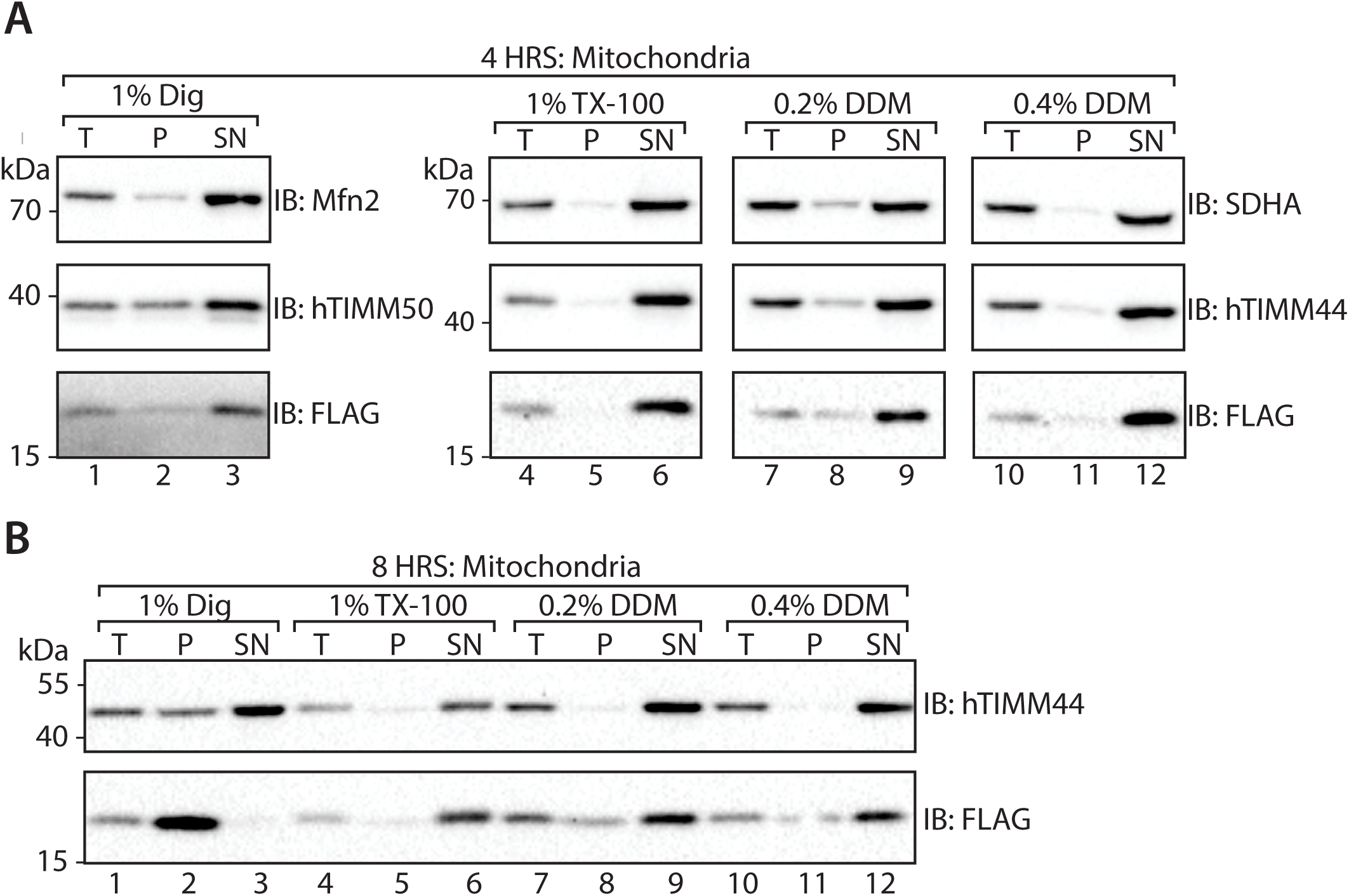
Solubility of MceC_3XFLAG_. Mitochondria were isolated from HEK293MceC- 3XFLAG cells induced with tetracycline (1 σg/mL) for (A) 4 HRS or (B) 8 HRS, were solubilised in digitonin (1% (w/v)), TX-100 (1% (v/v)) or DDM (0.2% and 0.4% (w/v)). Solubilised samples were separated by centrifugation and the total (T), pellet (P) and supernatant (SN) fractions analysed by SDS-PAGE and immunoblotting with the indicated antibodies.

**Supplemental Figure 2:**
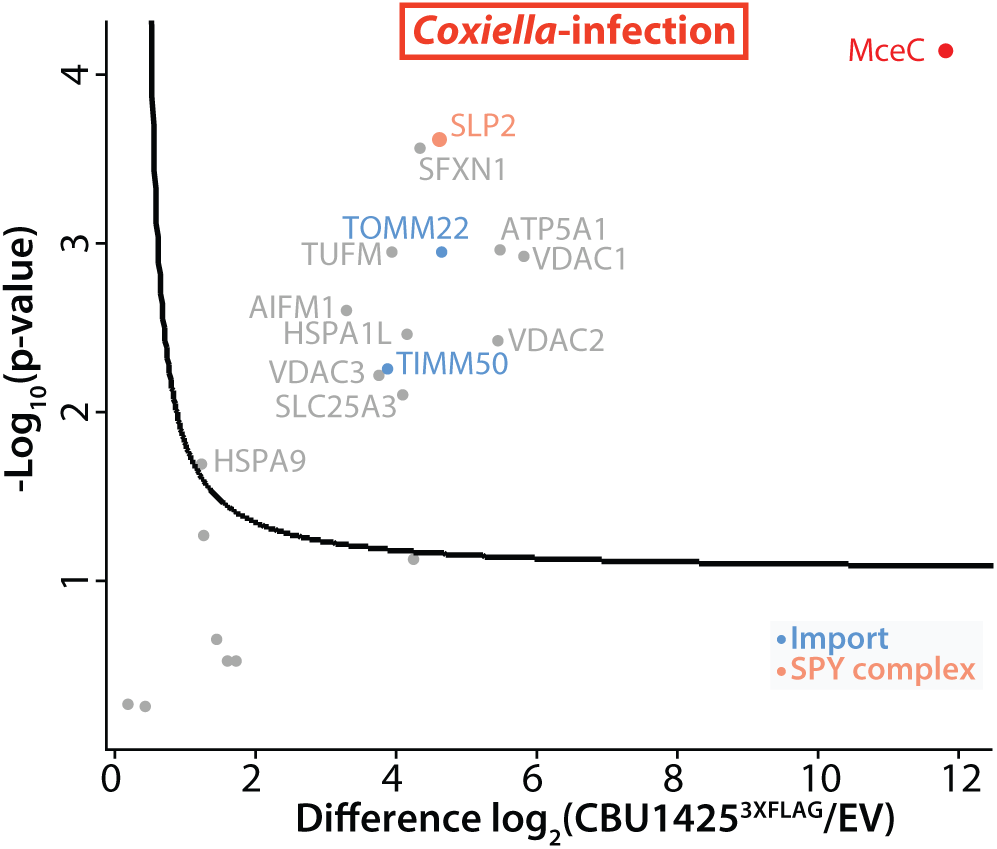
MceC immunoprecipitates resident mitochondrial proteins during *C. burnetii* infection. Mitochondria isolated from *C. bumetii-infected* control and HEK293 cells expressing MceC_3XFLAG_ for 4-hours were solubilised in digitonin (1% (w/v))-containing buffer and lysates subjected to anti-FLAG immunoprecipitation. Volcano plot showing proteins enriched following MceC immunoprecipitation as outlined in (A) compared to control ‘empty vector’ (EV) cells. Mitochondrial proteins plotted and each circle represents one protein. Gene names of selected proteins are used for labels. Horizontal axis shows the Log_2_(fold change) of MceC interacting partners and vertical axis shows -Log_10_(β-value) of two-sample Students T-test (FDR: 0.01 and s0: 1). n = 3 biological replicates. QC: Quality Control, IM: inner membrane. Colours correspond to key.

## Abbreviations

*C. burnetii*: *Coxiella burnetii*
CCV: Coxiella-containing vacuole Da: Dalton
IMS: intermembrane space
IMM: inner mitochondrial membrane
*L. pneumophila*: *Legionella pneumophila*
LFQ: label-free quantitation
Mce: Mitochondrial *Coxiella* effector protein
MTS: mitochondrial targeting signal
OMM: outer mitochondrial membrane
PK: proteinase K
T4SS: Type IV Secretion System
TIM: Translocase of the inner mitochondrial membrane
THP-1 cells: human monocyte derived macrophage cell line
TOM: translocase of the outer mitochondrial membrane
OXPHOS: oxidative phosphorylation

## References

1. T. R. Costa et al., Secretion systems in Gram-negative bacteria: structural and mechanistic insights. Nat Rev Microbiol 13, 343–359 (2015).

2. E. R. Green, J. Mecsas, Bacterial Secretion Systems: An Overview. Microbiol Spectr 4, (2016).

3. L. Gomez-Valero et al., More than 18,000 effectors in the Legionella genus genome provide multiple, independent combinations for replication in human cells. Proc Natl Acad Sci U S A 116, 2265–2273 (2019).

4. J. M. Park, S. Ghosh, T. J. O’Connor, Combinatorial selection in amoebal hosts drives the evolution of the human pathogen Legionella pneumophila. Nat Microbiol 5, 599–609 (2020).

5. A. Luhrmann, H. J. Newton, M. Bonazzi, Beginning to Understand the Role of the Type IV Secretion System Effector Proteins in Coxiella burnetii Pathogenesis. Curr Top Microbiol Immunol 413, 243–268 (2017).

6. M. Chien et al., The genomic sequence of the accidental pathogen Legionella pneumophila. Science 305, 1966–1968 (2004).

7. I. D. Hay, M. J. Belousoff, T. Lithgow, Structural Basis of Type 2 Secretion System Engagement between the Inner and Outer Bacterial Membranes. MBio 8, (2017).

8. D. Ghosal, Y. W. Chang, K. C. Jeong, J. P. Vogel, G. J. Jensen, In situ structure of the Legionella Dot/Icm type IV secretion system by electron cryotomography. EMBO Rep 18, 726–732 (2017).

9. D. Ghosal et al., Molecular architecture, polar targeting and biogenesis of the Legionella Dot/Icm T4SS. Nat Microbiol 4, 1173–1182 (2019).

10 L. J. Worrall et al., Near-atomic-resolution cryo-EM analysis of the Salmonella T3S injectisome basal body. Nature 540, 597–601 (2016).

11 M. L. Urbanus et al., Diverse mechanisms of metaeffector activity in an intracellular bacterial pathogen, Legionella pneumophila. Mol Syst Biol 12, 893 (2016).

12 P. M. Schneeberger, C. Wintenberger, W. van der Hoek, J. P. Stahl, Q fever in the Netherlands - 2007-2010: what we learned from the largest outbreak ever. Med Mal Infect 44, 339–353 (2014).

13 N. Arricau-Bouvery, A. Rodolakis, Is Q fever an emerging or re-emerging zoonosis? Vet Res 36, 327–349 (2005).

14 M. Maurin, D. Raoult, Q fever. Clin Microbiol Rev 12, 518–553 (1999).

15 D. Raoult, T. Marrie, J. Mege, Natural history and pathophysiology of Q fever. Lancet Infect Dis 5, 219–226 (2005).

16 C. C. Wielders et al., Characteristics of hospitalized acute Q fever patients during a large epidemic, The Netherlands. PLoS One 9, e91764 (2014).

17 J. G. Graham et al., Virulent Coxiella burnetii pathotypes productively infect primary human alveolar macrophages. Cell Microbiol 15, 1012–1025 (2013).

18 P. A. Beare et al., Dot/Icm type IVB secretion system requirements for Coxiella burnetii growth in human macrophages. MBio 2, e00175–00111 (2011).

19 K. L. Carey, H. J. Newton, A. Luhrmann, C. R. Roy, The Coxiella burnetii Dot/Icm system delivers a unique repertoire of type IV effectors into host cells and is required for intracellular replication. PLoS Pathog 7, e1002056 (2011).

20 J. H. Moffatt, P. Newton, H. J. Newton, Coxiella burnetii: turning hostility into a home. Cell Microbiol 17, 621–631 (2015).

21 C. L. Larson et al., Right on Q: genetics begin to unravel Coxiella burnetii host cell interactions. Future Microbiol 11, 919–939 (2016).

22 A. Omsland et al., Host cell-free growth of the Q fever bacterium Coxiella burnetii. Proc Natl Acad Sci U S A 106, 4430–4434 (2009).

23 P. A. Beare, C. L. Larson, S. D. Gilk, R. A. Heinzen, Two systems for targeted gene deletion in Coxiella burnetii. Appl Environ Microbiol 78, 4580–4589 (2012).

24 C. Chen et al., Large-scale identification and translocation of type IV secretion substrates by Coxiella burnetii. Proc Natl Acad Sci U S A 107, 21755–21760 (2010).

25 D. E. Voth et al., The Coxiella burnetii cryptic plasmid is enriched in genes encoding type IV secretion system substrates. J Bacteriol 193, 1493–1503 (2011).

26 P. A. Beare et al., Essential role for the response regulator PmrA in Coxiella burnetii type 4B secretion and colonization of mammalian host cells. J Bacteriol 196, 1925–1940 (2014).

27 K. M. Sandoz, P. A. Beare, D. C. Cockrell, R. A. Heinzen, Complementation of Arginine Auxotrophy for Genetic Transformation of Coxiella burnetii by Use of a Defined Axenic Medium. Appl Environ Microbiol 82, 3042–3051 (2016).

28 L. F. Fielden et al., A Farnesylated Coxiella burnetii Effector Forms a Multimeric Complex at the Mitochondrial Outer Membrane during Infection. Infect Immun 85, (2017).

29 A. J. Anderson, T. D. Jackson, D. A. Stroud, D. Stojanovski, Mitochondria-hubs for regulating cellular biochemistry: emerging concepts and networks. Open Biol 9, 190126 (2019).

30 N. Pfanner, B. Warscheid, N. Wiedemann, Mitochondrial proteins: from biogenesis to functional networks. Nat Rev Mol Cell Biol 20, 267–284 (2019).

31 E. L. Mills, B. Kelly, L. A. J. O’Neill, Mitochondria are the powerhouses of immunity. Nat Immunol 18, 488–498 (2017).

32 L. F. Fielden, Y. Kang, H. J. Newton, D. Stojanovski, Targeting mitochondria: how intravacuolar bacterial pathogens manipulate mitochondria. Cell Tissue Res 367, 141–154 (2017).

33 A. Spier, F. Stavru, P. Cossart, Interaction between Intracellular Bacterial Pathogens and Host Cell Mitochondria. Microbiol Spectr 7, (2019).

34 V. Tiku, M. W. Tan, I. Dikic, Mitochondrial Functions in Infection and Immunity. Trends Cell Biol 30, 263–275 (2020).

35 P. Escoll et al., Legionella pneumophila Modulates Mitochondrial Dynamics to Trigger Metabolic Repurposing of Infected Macrophages. Cell Host Microbe 22, 302–316 e307 (2017).

36 A. Luhrmann, C. R. Roy, Coxiella burnetii inhibits activation of host cell apoptosis through a mechanism that involves preventing cytochrome c release from mitochondria. Infect Immun 75, 5282–5289 (2007).

37 L. Klingenbeck, R. A. Eckart, C. Berens, A. Luhrmann, The Coxiella burnetii type IV secretion system substrate CaeB inhibits intrinsic apoptosis at the mitochondrial level. Cell Microbiol 15, 675–687 (2013).

38 A. Luhrmann, C. V. Nogueira, K. L. Carey, C. R. Roy, Inhibition of pathogen-induced apoptosis by a Coxiella burnetii type IV effector protein. Proc Natl Acad Sci U S A 107, 18997–19001 (2010).

39 A. Omsland et al., Isolation from animal tissue and genetic transformation of Coxiella burnetii are facilitated by an improved axenic growth medium. Appl Environ Microbiol 77, 3720–3725 (2011).

40 Y. Kang et al., Tim29 is a novel subunit of the human TIM22 translocase and is involved in complex assembly and stability. Elife 5, (2016).

41 K. Jaton, O. Peter, D. Raoult, J. D. Tissot, G. Greub, Development of a high throughput PCR to detect Coxiella burnetii and its application in a diagnostic laboratory over a 7-year period. New Microbes New Infect 1, 6–12 (2013).

42 A. J. Johnston et al., Insertion and assembly of human tom7 into the preprotein translocase complex of the outer mitochondrial membrane. J Biol Chem 277, 42197–42204 (2002).

43 J. M. Graham, Purification of a crude mitochondrial fraction by density-gradient centrifugation. Curr Protoc Cell Biol Chapter 3, Unit 3 4 (2001).

44 H. Schagger, G. von Jagow, Tricine-sodium dodecyl sulfate-polyacrylamide gel electrophoresis for the separation of proteins in the range from 1 to 100 kDa. Anal Biochem 166, 368–379 (1987).

45 H. Schagger, Tricine-SDS-PAGE. Nat Protoc 1, 16–22 (2006).

46 Y. Kang et al., Function of hTim8a in complex IV assembly in neuronal cells provides insight into pathomechanism underlying Mohr-Tranebjaerg syndrome. Elife 8, (2019).

47 Y. Kang et al., Sengers Syndrome-Associated Mitochondrial Acylglycerol Kinase Is a Subunit of the Human TIM22 Protein Import Complex. Mol Cell 67, 457–470 e455 (2017).

48 J. Rappsilber, Y. Ishihama, M. Mann, Stop and go extraction tips for matrix-assisted laser desorption/ionization, nanoelectrospray, and LC/MS sample pretreatment in proteomics. Anal Chem 75, 663–670 (2003).

49 J. Rappsilber, M. Mann, Y. Ishihama, Protocol for micro-purification, enrichment, pre-fractionation and storage of peptides for proteomics using StageTips. Nat Protoc 2, 1896–1906 (2007).

50 J. Cox, M. Mann, MaxQuant enables high peptide identification rates, individualized p.p.b.- range mass accuracies and proteome-wide protein quantification. Nat Biotechnol 26, 1367–1372 (2008).

51 J. Cox et al., Accurate proteome-wide label-free quantification by delayed normalization and maximal peptide ratio extraction, termed MaxLFQ. Mol Cell Proteomics 13, 2513–2526 (2014).

52 S. Tyanova et al., The Perseus computational platform for comprehensive analysis of (prote)omics data. Nat Methods 13, 731–740 (2016).

53 S. E. Calvo, K. R. Clauser, V. K. Mootha, MitoCarta2.0: an updated inventory of mammalian mitochondrial proteins. Nucleic Acids Res 44, D1251–1257 (2016).

54 D. J. Pagliarini et al., A mitochondrial protein compendium elucidates complex I disease biology. Cell 134, 112–123 (2008).

55 E. W. Deutsch et al., The ProteomeXchange consortium in 2020: enabling ‘big data’ approaches in proteomics. Nucleic Acids Res 48, D1145–D1152 (2020).

56 Y. Perez-Riverol et al., The PRIDE database and related tools and resources in 2019: improving support for quantification data. Nucleic Acids Res 47, D442–D450 (2019).

57 C. Noroy, T. Lefrancois, D. F. Meyer, Searching algorithm for Type IV effector proteins (S4TE) 2.0: Improved tools for Type IV effector prediction, analysis and comparison in proteobacteria. PLoS Comput Biol 15, e1006847 (2019).

58 M. Lazarou et al., Inhibition of Bak activation by VDAC2 is dependent on the Bak transmembrane anchor. J Biol Chem 285, 36876–36883 (2010).

59 L. E. Formosa et al., Characterization of mitochondrial FOXRED1 in the assembly of respiratory chain complex I. Hum Mol Genet 24, 2952–2965 (2015).

60 M. Lazarou, M. McKenzie, A. Ohtake, D. R. Thorburn, M. T. Ryan, Analysis of the assembly profiles for mitochondrial- and nuclear-DNA-encoded subunits into complex I. Mol Cell Biol 27, 4228–4237 (2007).

61 C. S. Palmer et al., MiD49 and MiD51, new components of the mitochondrial fission machinery. EMBO Rep 12, 565–573 (2011).

62 D. H. Hock et al., HIGD2A is required for assembly of the COX3 module of human mitochondrial complex IV. Mol Cell Proteomics, (2020).

63 B. Padmanabhan et al., Biogenesis of the spacious Coxiella-containing vacuole depends on host transcription factors TFEB and TFE3. Infect Immun, (2019).

64 N. R. Sims, M. F. Anderson, Isolation of mitochondria from rat brain using Percoll density gradient centrifugation. Nat Protoc 3, 1228–1239 (2008).

65 M. R. Wieckowski, C. Giorgi, M. Lebiedzinska, J. Duszynski, P. Pinton, Isolation of mitochondria-associated membranes and mitochondria from animal tissues and cells. Nat Protoc 4, 1582–1590 (2009).

66 D. A. Clayton, G. S. Shadel, Purification of mitochondria by sucrose step density gradient centrifugation. Cold Spring Harb Protoc 2014, pdb prot080028 (2014).

67 H. T. Hornig-Do et al., Isolation of functional pure mitochondria by superparamagnetic microbeads. Anal Biochem 389, 1–5 (2009).

68 M. M. Weber et al., Identification of Coxiella burnetii type IV secretion substrates required for intracellular replication and Coxiella-containing vacuole formation. J Bacteriol 195, 3914–3924 (2013).

69 C. L. Larson et al., Coxiella burnetii effector proteins that localize to the parasitophorous vacuole membrane promote intracellular replication. Infect Immun 83, 661–670 (2015).

70 P. A. Beare et al., Comparative genomics reveal extensive transposon-mediated genomic plasticity and diversity among potential effector proteins within the genus Coxiella. Infect Immun 77, 642–656 (2009).

71 E. J. van Schaik, E. D. Case, E. Martinez, M. Bonazzi, J. E. Samuel, The SCID Mouse Model for Identifying Virulence Determinants in Coxiella burnetii. Front Cell Infect Microbiol 7, 25 (2017).

72 M. Liu, G. M. Conover, R. R. Isberg, Legionella pneumophila EnhC is required for efficient replication in tumour necrosis factor alpha-stimulated macrophages. Cell Microbiol 10, 1906–1923 (2008).

73 M. Liu et al., The Legionella pneumophila EnhC protein interferes with immunostimulatory muramyl peptide production to evade innate immunity. Cell Host Microbe 12, 166–176 (2012).

74 Y. Kang, L. F. Fielden, D. Stojanovski, Mitochondrial protein transport in health and disease. Semin Cell Dev Biol 76, 142–153 (2018).

75 M. G. Teese, D. Langosch, Role of GxxxG Motifs in Transmembrane Domain Interactions. Biochemistry 54, 5125–5135 (2015).

76 Y. Ohba, T. MacVicar, T. Langer, Regulation of mitochondrial plasticity by the i-AAA protease YME1L. Biol Chem, (2020).

77 T. M. Greco, M. A. Kennedy, I. M. Cristea, Proteomic Technologies for Deciphering Local and Global Protein Interactions. Trends Biochem Sci 45, 454–455 (2020).

78 Z. Lifshitz etal., Identification of novel Coxiella burnetii Icm/Dot effectors and genetic analysis of their involvement in modulating a mitogen-activated protein kinase pathway. Infect Immun 82, 3740–3752 (2014).

79 J. Schmolders et al., Comparative Proteomics of Purified Pathogen Vacuoles Correlates Intracellular Replication of Legionella pneumophila with the Small GTPase Ras-related protein 1 (Rap1). Mol Cell Proteomics 16, 622–641 (2017).

80 J. Naujoks et al., IFNs Modify the Proteome of Legionella-Containing Vacuoles and Restrict Infection Via IRG1-Derived Itaconic Acid. PLoS Pathog 12, e1005408 (2016).

81 P. M. Jean Beltran, R. A. Mathias, I. M. Cristea, A Portrait of the Human Organelle Proteome In Space and Time during Cytomegalovirus Infection. Cell Syst 3, 361–373 e366 (2016).

82 Y. Hashimoto, X. Sheng, L. A. Murray-Nerger, I. M. Cristea, Temporal dynamics of protein complex formation and dissociation during human cytomegalovirus infection. Nat Commun 11, 806 (2020).

83 K. C. Cook, I. M. Cristea, Location is everything: protein translocations as a viral infection strategy. Curr Opin Chem Biol 48, 34–43 (2019).

84 W. Schafer et al., Nuclear trafficking of the anti-apoptotic Coxiella burnetii effector protein AnkG requires binding to p32 and Importin-alpha1. Cell Microbiol 19, (2017).

85 V. Hung et al., Proteomic mapping of cytosol-facing outer mitochondrial and ER membranes in living human cells by proximity biotinylation. Elife 6, (2017).

86 H. W. Rhee et al., Proteomic mapping of mitochondria in living cells via spatially restricted enzymatic tagging. Science 339, 1328–1331 (2013).

87 K. A. T. Makepeace et al., Improving Identification of In-organello Protein-Protein Interactions Using an Affinity-enrichable, Isotopically Coded, and Mass Spectrometry-cleavable Chemical Crosslinker. Mol Cell Proteomics 19, 624–639 (2020).

88 A. Mahdavi et al., Identification of secreted bacterial proteins by noncanonical amino acid tagging. Proc Natl Acad Sci U S A 111,433–>438 (2014).

89 A. J. Link et al., Discovery of aminoacyl-tRNA synthetase activity through cell-surface display of noncanonical amino acids. Proc Natl Acad Sci U S A 103, 10180–10185 (2006).

90 A. Mahdavi et al., Engineered Aminoacyl-tRNA Synthetase for Cell-Selective Analysis of Mammalian Protein Synthesis. J Am Chem Soc 138, 4278–4281 (2016).

91 P. Escoll, S. Mondino, M. Rolando, C. Buchrieser, Targeting of host organelles by pathogenic bacteria: a sophisticated subversion strategy. Nat Rev Microbiol 14, 5–19 (2016).

92 G. Lutfullahoglu-Bal, A. Keskin, A. B. Seferoglu, C. D. Dunn, Bacterial tail anchors can target to the mitochondrial outer membrane. Biol Direct 12, 16 (2017).

93 J. E. Muller et al., Mitochondria can recognize and assemble fragments of a beta-barrel structure. Mol Biol Cell 22, 1638–1647 (2011).

94 C. Ott, M. Utech, M. Goetz, T. Rudel, V. Kozjak-Pavlovic, Requirements for the import of neisserial Omp85 into the outer membrane of human mitochondria. Biosci Rep 33, e00028 (2013).

95 V. Kozjak-Pavlovic et al., Bacterial porin disrupts mitochondrial membrane potential and sensitizes host cells to apoptosis. PLoS Pathog 5, e1000629 (2009).

96 V. Kozjak-Pavlovic, C. Ott, M. Gotz, T. Rudel, Neisserial Omp85 protein is selectively recognized and assembled into functional complexes in the outer membrane of human mitochondria. J Biol Chem 286, 27019–27026 (2011).

97 H. Niu, V. Kozjak-Pavlovic, T. Rudel, Y. Rikihisa, Anaplasma phagocytophilum Ats-1 is imported into host cell mitochondria and interferes with apoptosis induction. PLoS Pathog 6, e1000774 (2010).

98 S. R. Shames, M. A. Croxen, W. Deng, B. B. Finlay, The type III system-secreted effector EspZ localizes to host mitochondria and interacts with the translocase of inner mitochondrial membrane 17b. Infect Immun 79, 4784–4790 (2011).

99. A. Muller et al., Targeting of the pro-apoptotic VDAC-like porin (PorB) of Neisseria gonorrhoeae to mitochondria of infected cells. EMBO J 19, 5332–5343 (2000).

100. S. Kutik et al., Dissecting membrane insertion of mitochondrial beta-barrel proteins. Cell 132, 1011–1024 (2008).

101 R. Gomkale et al., Defining the Substrate Spectrum of the TIM22 Complex Identifies Pyruvate Carrier Subunits as Unconventional Cargos. Curr Biol 30, 1119–1127 e1115 (2020).

102 S. Ahola, T. Langer, T. MacVicar, Mitochondrial Proteolysis and Metabolic Control. Cold Spring Harb Perspect Biol 11, (2019).

103 M. J. Baker, T. Tatsuta, T. Langer, Quality control of mitochondrial proteostasis. Cold Spring Harb Perspect Biol 3, (2011).

104 T. Wai et al., The membrane scaffold SLP2 anchors a proteolytic hub in mitochondria containing PARL and the i-AAA protease YME1L. EMBO Rep 17, 1844–1856 (2016).

105 T. MacVicar et al., Lipid signalling drives proteolytic rewiring of mitochondria by YME1L. Nature 575, 361–365 (2019).

106 R. Anand et al., The i-AAA protease YME1L and OMA1 cleave OPA1 to balance mitochondrial fusion and fission. J Cell Biol 204, 919–929 (2014).

107 M. J. Baker et al., Stress-induced OMA1 activation and autocatalytic turnover regulate OPA1- dependent mitochondrial dynamics. EMBO J 33, 578–593 (2014).

108 P. Mishra, V. Carelli, G. Manfredi, D. C. Chan, Proteolytic cleavage of Opa1 stimulates mitochondrial inner membrane fusion and couples fusion to oxidative phosphorylation. Cell Metab 19, 630–641 (2014).

109 M. Suzuki, O. Danilchanka, J. J. Mekalanos, Vibrio cholerae T3SS effector VopE modulates mitochondrial dynamics and innate immune signaling by targeting Miro GTPases. Cell Host Microbe 16, 581–591 (2014).

110 F. Stavru, F. Bouillaud, A. Sartori, D. Ricquier, P. Cossart, Listeria monocytogenes transiently alters mitochondrial dynamics during infection. Proc Natl Acad Sci U S A 108, 3612–3617 (2011).

111 F. Stavru, A. E. Palmer, C. Wang, R. J. Youle, P. Cossart, Atypical mitochondrial fission upon bacterial infection. Proc Natl Acad Sci U S A 110, 16003–16008 (2013).

